# Flexible perception of face attributes under naturalistic visual constraints

**DOI:** 10.64898/2026.01.26.701887

**Authors:** Shiyin Li, You Wu, Xiaoqian Yan

## Abstract

How does human brain adapt face perception to naturalistic visual constraints? While faces are typically perceived under dynamic and uncertain conditions, most studies use static presentation and enough temporal exposure, leaving the neural mechanisms of adaptive face processing poorly understood. Here, we employed steady-state evoked potential electroencephalography (EEG) to track how perception of multiple face attributes (age, emotion, gender, and race) adjusts to two ecologically relevant scenarios: (1) faces gradually sharpening from blur to clear (mimicking approaching from a distance), and (2) faces presented with incrementally increasing exposure times (mimicking brief, flashed encounters). By progressively increasing sensory input (via spatial frequency content in Experiment 1 and exposure time in Experiment 2) in a stepwise manner, we show that emotion and race categorization emerge early, even under high blur (e.g., 4.89 cycles/image) or brief exposures (e.g., 41.7-50 ms), reflecting coarse visual processing. In contrast, age discrimination requires higher clarity (e.g., 7.31 cycles/image) but shorter exposure (e.g., 41.7 ms) if clear images are presented. In both scenarios, gender processing exhibits the strongest dependence on clarity (e.g., 10.94 cycles/image) and time (e.g., 66.7 ms). Representational similarity analyses further show that reliable response patterns for emotion and race emerge earlier in the posterior brain region relative to those for gender in both experiments. Together, these results identify a flexible temporal order for face perception that adapts to naturalistic visual constraints, bridging the gap between controlled laboratory paradigms and naturalistic social vision.

## Introduction

Human face perception operates under constantly changing visual conditions—whether recognizing a distant friend approaching or interpreting a stranger’s expression in a fleeting glance. Yet, neuroscientific studies overwhelmingly rely on static, high-clarity faces, neglecting how the brain adapts perception to real-world constraints like motion, blur, or time pressure. Here, we address this gap by investigating how perception of “visually derived face attributes” (e.g. age, emotion, gender, and race) dynamically adjusts to two naturalistic scenarios: (1) progressive clarity (blur-to-clear transitions, mimicking approaching faces) and (2) increasing exposure time (brief-to-prolonged presentations, mimicking flashed encounters).

Faces convey rich social signals—age, emotion, gender, identity, and more ^1^. While identity-related processing is restricted to familiar faces, other features can be extracted universally. Recent neuroimaging advances reveal a hierarchical temporal dynamic in face processing: some visually derived attributes such as age and emotion are decoded earlier than identity, supporting a coarse-to-fine processing model ^2–7^. For instance, ^3^ reported that age and gender information emerged early in MEG representations (∼ 60-70 ms), preceding identity decoding by ∼ 20 ms. Similarly, ^5^ observed earlier extraction of age and emotion information (∼ 110 ms) compared to identity (∼ 235 ms). These findings suggest prioritized extraction of socially salient traits, possibly reflecting adaptive perceptual mechanisms for rapid social evaluation. However, four key limitations remain unresolved in characterizing non-identity face representations: (1) while some studies report early gender decoding (e.g., ^3^), others show weaker effects (e.g., ^5^), suggesting potential methodological or stimulus-dependent variability; (2) the temporal dynamics of race perception remain unexplored in this framework, despite its fundamental role in social cognition; (3) no study has systematically examined the neural temporal processing order across all four key social attributes (age, emotion, gender, and race) within a unified experimental paradigm; (4) while existing work employs high-resolution face stimuli with prolonged exposure durations (≥ 200 ms) to ensure robust neural representations, it cannot reveal the adaptive mechanisms underlying face perception under ecologically relevant uncertainty.

In the current study, we implemented a sweep steady-state visual evoked potential (SSVEP) paradigm to track implicit neural measures of categorization of four face attributes, i.e., age, emotion, gender, and race. We constrained sensory input by parametrically varying either image spatial frequency content (i.e., spatial resolution, Experiment 1), or image exposure time (i.e., presentation duration, Experiment 2) ^8–10^. By measuring the effects of increasing spatial frequency/exposure time on implicit neural indices of face attribute categorization, we identified the minimal amount of sensory input required to elicit each brain function under conditions that mimic the dynamic nature of real-world perception. We also used representational similarity analysis (RSA) to investigate how reliable neural representations to distinct face attributes evolve over time. Our results uncover a spatiotemporally dissociable neural patterns for adaptive face perception. We demonstrate that emotion and race categorization emerge early, even under high blur or brief exposures, reflecting reliance on coarse visual processing. In contrast, age discrimination follows a distinct processing profile: it requires higher visual clarity but achieves reliable representation with relatively short exposures when visual input is unobscured. In both scenarios, gender processing exhibits the strongest dependence on clarity and time. These findings advance our understanding of naturalistic social cognition and offer new avenues for studying clinical populations (e.g., autism, prosopagnosia) with atypical face perception under uncertainty.

## Materials and methods

### Participants

Forty-four participants from Fudan University were recruited to take part in our experiments (24 females, mean age 22.1 ± 2.2 years). Data from seven participants were excluded due to excessive noise/muscular artifacts during EEG recording. The final sample consisted of thirty-eight participants: sixteen (8 females, mean age 22.3 ± 2.6 years) in Experiment 1, ten (6 females, mean age 21.9 ± 1.4 years) in Experiment 2-1, and sixteen (7 females, mean age 22.2 ± 1.6 years) in Experiment 2-2. All participants reported normal or corrected-to-normal visual acuity. They were all right-handed according to self-report, with normal or corrected-to-normal vision. Written consent was obtained prior to test for a study approved by the Ethics Committee of Fudan University (NO. FE232451).

### Stimuli

Face images selected from Tsinghua ^11^ and KDEF ^12^ dataset were used as stimuli. To measure the categorization responses in four face attributes, i.e., age, emotion, gender, and race, we selected 21 young Asian female neutral faces, 21 old Asian female neutral faces, 21 young Asian female happy faces, 21 young Asian female sad faces, 21 young Asian male neutral faces, and 21 young Caucasian female neutral faces from these two image sets. All face stimuli were cropped to keep only the head area with a height and width of 200 pixels and converted to grayscale. When viewed from a distance of 70 cm, each image subtended a visual angle of approximately 9.80°.

To study the modulation effect of spatial frequency (SF) on face categorization, in Experiment 1, we applied a low-pass gaussian filter to the images at 9 logarithmic cut-off steps ^10^. Before applying the gaussian filters, we first normalized the full-spectrum grayscale images to achieve a global luminance with a mean of 0 and a standard deviation (SD) of 1. Filtered images were then obtained by applying fast Fourier transforms to each image and multiplying the Fourier energy by Gaussian filters. The images were low-pass filtered at spatial frequencies ranging from 4 to 20 cycles per image (cpi) in 9 steps, i.e. 4, 4.89, 5.98, 7.31, 8.94, 10.94, 13.37, 16.36 and 20 cpi (Figure 1). This resulted in a total of 189 images across SF steps. The luminance and contrast of all face images from each SF step were adjusted to match the mean values of the original full-spectrum images to guarantee equal global luminance and contrast values both within and across SF steps. We did not apply further manipulations on the images. To rule out the possibility that salient low-level cues specific to some image categories drove early processing, we ran image spatial frequency analysis ^13^ and found that the spectral power distributions were very similar across categories (Supplementary Figure S1).

**Figure 1.**
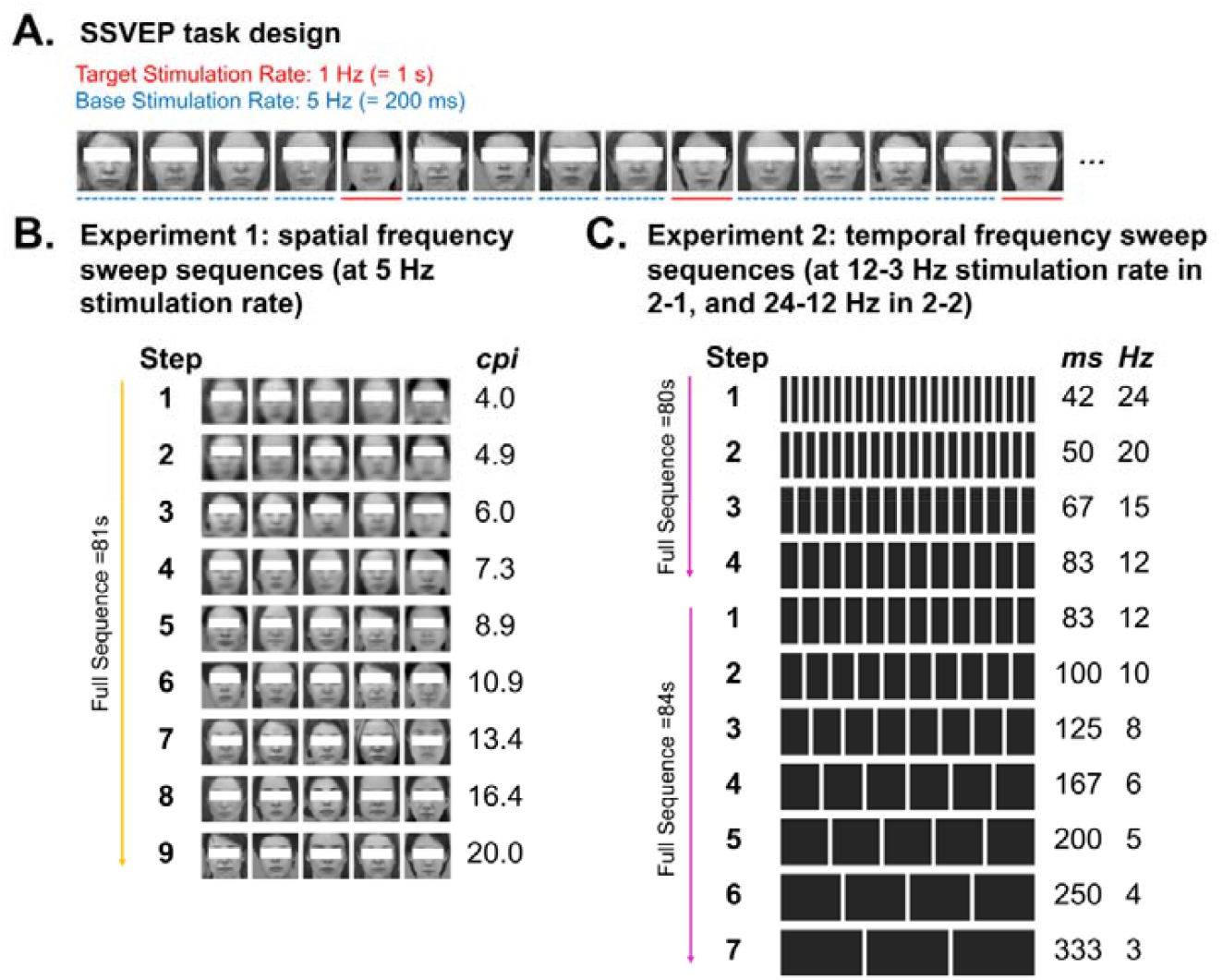
Design overview. **(A)**. Task design with low-pass filtered (20 cycles/image, cpi) images. In the example sequence, implicit race categorization response is measured. Young <u>Caucasian</u> female face images are presented at a fixed rate of 5 Hz (i.e., 200 ms per image), with young <u>Asian</u> female face images appearing at every 1 s. **(B)**. Sequences in Experiment 1. Images in this experiment are low-pass filtered with different spatial frequency contents (from 4 cpi to 20 cpi in 9 steps). C. Sequences in Experiment 2 (2-1 and 2-2). Images in the experiments are shown with different temporal frequency (from 24 Hz to 12 Hz in Experiment 2-2, and from 12 Hz to 3 Hz in Experiment 2-1).

### Procedure

#### Experiment 1: spatial frequency sweep SSVEP paradigm

To investigate the minimal spatial frequency required for successful face categorization across different face attributes, we employed a sweep SSVEP EEG paradigm ^8–10^. The experiment comprised four blocks corresponding to four face conditions: (1) in the age condition, <u>young</u> Asian female neutral faces were used as target images and <u>old</u> Asian female neutral faces were used as baseline stimuli; (2) in the emotion condition, young Asian female <u>sad</u> faces were used as target images and young Asian female <u>happy</u> faces were used as baseline images; (3) in the gender condition, young Asian <u>female</u> neutral faces were used as target images and young Asian <u>male</u> neutral faces were used as baseline stimuli; and (4) in the race condition, young <u>Asian</u> female neutral faces were used as target images and young <u>Caucasian</u> female neutral faces were used as baseline stimuli. During the experiment, block presentation order was randomized across participants. Each block contained 15 sequences. During each 81-s sequence, face images were presented at a fixed rate of 5 Hz (i.e., 5 images by second) with target stimuli inserted at every 5^th^ image (i.e., 1 Hz) (**Figure 1A**). During each sequence, a sweep design was used to systematically modulate the image SF contents. Specifically, the SF content of the images gradually increased every 9 s over the course of 9 sequential SF steps. In this way, the images initially appeared blurry (i.e., low-pass filtered), and gradually sharpened into clearer images over the course of 81 s (**Figure 1B**). In these four conditions, categorical responses at 1 Hz (and harmonics) could only be evoked and thus measured if participants could implicitly discriminate the differences between the rapidly represented two face image categories in a sequence.

During each stimulation sequence, the face images were presented at the center of the screen, with a blue fixation cross overlaid on top of the face images and centered on the screen as well. Participants were instructed to respond as quickly and as accurately as possible by pressing a space bar when the color of the fixation cross changed from blue to red (10 times, 200-ms duration each time), while at the same time being instructed to monitor the face images on the screen. The experiment was self-paced. Participants controlled the progression of the experiment by pressing the spacebar on a keyboard to proceed to the next trial or block. The entire experiment took approximately 120 min, including breaks. Behavioral data from one participant was excluded due to technical failures during data export. For the remaining 15 participants, the mean accuracy on the behavioral task was 90.5% (SE = 2.1%), with a mean correct response time (RT) of 503.9 ms (SE = 19.3 ms).

#### Experiment 2: temporal frequency sweep SSVEP paradigm

The task design of Experiment 2 was similar to previous studies using a sweep SSVEP EEG paradigm ^8,14^. Images from two categories were presented at varying durations in a stepwise manner. Two experiments were conducted to determine the minimal image exposure time required to measure rapid face attributes categorization.

In the pilot Experiment 2-1, stimuli were presented at frequencies of 12, 10, 8, 6, 5, 4, and 3 Hz, corresponding to individual image durations of 83, 100, 125, 167, 200, 250, and 333 ms, respectively. The experiment included four blocks, each representing a distinct face condition. All face images were identical to those used in Experiment 1 but with full-spectrum. Each block consisted of 14 sequences, with every 84 s sequence containing 7 contiguous 12 s steps of stimulus presentation (**Figure 1C**). These steps followed a fast-to-slow frequency sweep (from 12 Hz to 3 Hz with a fixed order). At each step, baseline images were presented at a fixed rate, with target face images embedded at 1 s intervals. Block order was randomized across participants. The entire experiment lasted approximately 120 min, including breaks. The results showed significant categorization responses across all four face conditions even when images were presented at 12 Hz (i.e., 83 ms per image) (full results are available in the supplementary file, section 3), indicating that even higher temporal frequency ranges were required to reach the perceptual threshold for rapid face categorization.

In Experiment 2-2, we increased the maximum image presentation frequency to 24 Hz. The experiment consisted of four blocks, each containing 12 sequences. Each 80 s sequence comprised four contiguous 20 s steps of stimulus presentation at frequencies of 24, 20, 15, and 12 Hz, following a descending frequency sweep (**Figure 1C**). The entire experiment lasted approximately 90 minutes, including breaks. Participants performed the same task as in Experiment 1, with a mean response accuracy of 92.9% (SE = 1.8%), and a mean correct RT of 492.6 ms (SE = 18.8 ms).

#### EEG acquisition

EEG signals were recorded using a 64-electrode EEG acquisition system (actiCAP, Brain Products GmbH, Gilching, Germany) configured according to the international 10-20 electrode placement protocol. Neural activity was digitized at a sampling rate of 1 kHz, with Cz designated as the online reference electrode. Electrode-scalp contact integrity was maintained by filling each electrode with conductive gel and adjusting impedances to remain below 5 kΩ for all electrodes throughout the experiment. Data acquisition was managed using BrainVision Recorder software (Version 1.21, Brain Products GmbH, Gilching, Germany).

### Analysis

#### EEG Preprocessing

EEG data analysis was performed using the software Letswave6 (http://www.nocions.org/letswave), running on MATLAB R2022a (MathWorks, USA). The EEG data were initially band-pass filtered at 0.01-100 Hz using a fourth-order zero-phase Butterworth filter, followed by down-sampling to 250 Hz to reduce computational load. The data sequence was then segmented relative to the starting trigger of each trial, with a duration of 81 s in Experiment 1, 84 s in Experiment 2-1, and 80 s in Experiment 2-2. To correct for eye blinks, independent component analysis (ICA) was applied to all participants who showed obvious blinking components. Individual electrode with artifacts were interpolated using their three neighboring electrodes, with no interpolated electrodes per participant for Experiment 1, 0.1 ± 0.3 for Experiment 2-1 and 0.38 ± 0.7 for Experiment 2-2. The cleaned data were then referenced to the average of all 63 electrodes (excluding the online reference Cz electrode).

#### EEG Frequency Domain Analysis

For each face condition, the preprocessed EEG data were cropped into epochs according to each SF or temporal frequency (TF) step (Experiment 1 = 9 × 9 s epochs, Experiment 2-1 = 7 × 12 s epochs, Experiment 2-2 = 4 × 20 s epochs). A Fast Fourier Transform (FFT) was applied to each averaged SF/TF step epoch and the amplitude spectra were extracted, with a frequency resolution of 0.111 Hz (1/9 s) for Experiment1, 0.083 Hz (1/12 s) for Experiment 2-1 and 0.05 Hz (1/20 s) for Experiment 2-2. To correct for variations in baseline noise level around each frequency of interest, two methods were used: (1) the mean amplitude of the neighboring 8 bins (4 by each side) were calculated and divided from each target frequency bin to display EEG spectra in signal-to-noise ratio (SNR), allowing better visualization of small responses (only for **Figure 2, and Supplementary Figure S4**) ^10,14,15^ and (2) subtraction of the EEG noise from target bins (baseline subtraction) to quantify responses in microvolt. At each SF/TF step, the response was quantified with a summation of response at 1 Hz and harmonics up to 8 Hz for Experiment 2-1, Experiment 2-2 and 9 Hz for Experiment 1 (excluding the frequency that coincided with the baseline stimuli presentation frequency according to every step) ^10,14,15^. The number of harmonics selected in the analyses was determined according to the grand-averaged response patterns across all participants, all electrodes and across face conditions ^15,16^.

**Figure 2.**
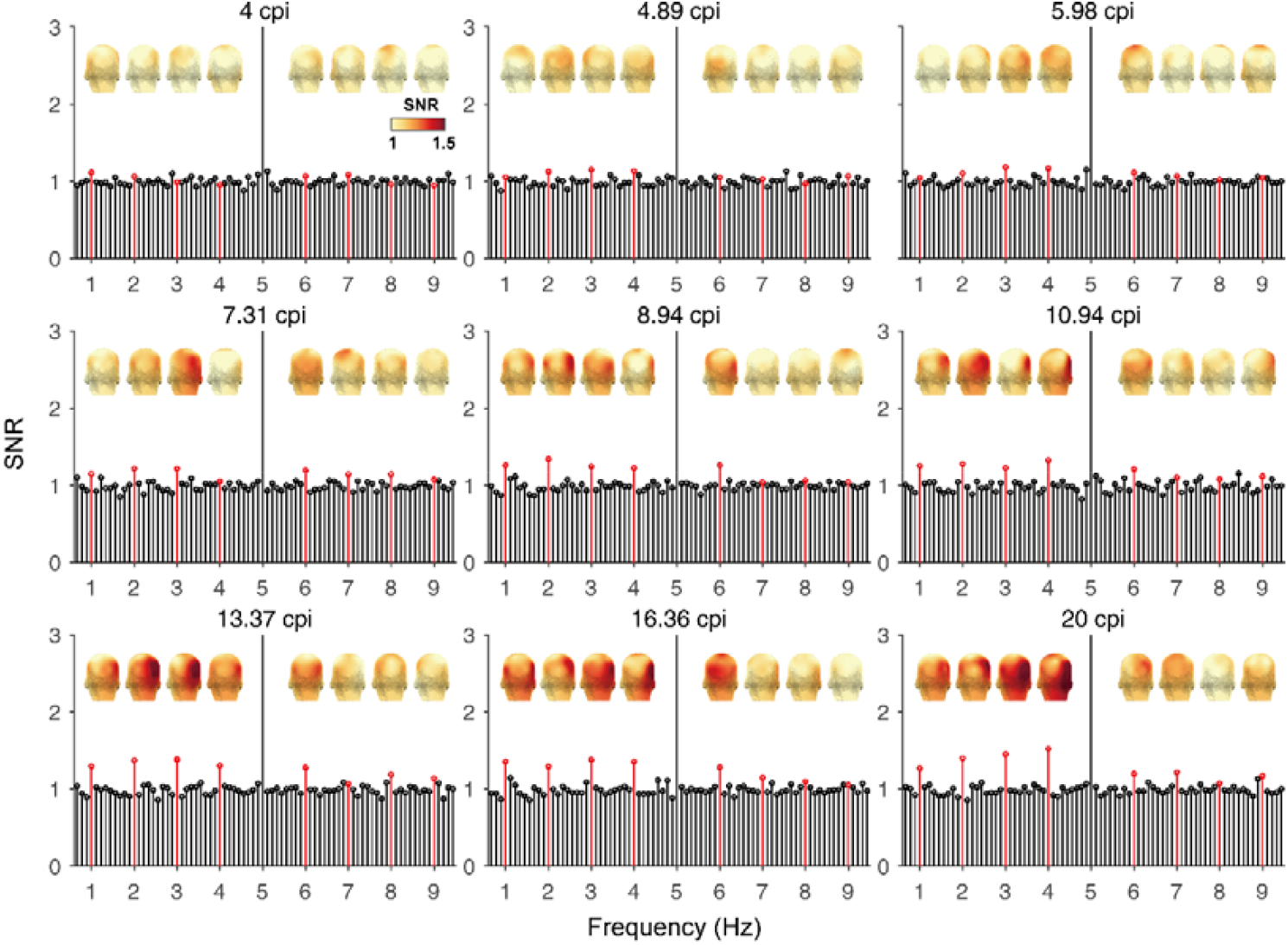
Grand-averaged frequency spectra in signal-to-noise ratio (SNR) across four face conditions as a function of image SF content over the OT ROI. The frequency bins at every 1 Hz step (up to 9 Hz, excluding the base image presentation rate of 5 Hz) are highlighted in red lines. Three-D topographies of 1 Hz and its selected harmonics are shown above.

In both experiments, based on the mean response amplitude distribution across all conditions and participants, we selected two regions of interest (ROIs) for further analyses: (1) right occipitotemporal (rOT) ROI (P6, P8, and PO8), and (2) left occipitotemporal (lOT) ROI (P5, P7, and PO7), to quantify the target categorization responses at 1 Hz and harmonics.

To ensure that the experimental noise level in the EEG signal did not vary across different face conditions, we measured the amplitude of noise bins in frequencies up to 9 Hz excluding the image presentation frequency (e.g., 5 Hz in Experiment 1) and the target categorization frequencies (1 Hz and harmonics) in the OT ROIs. We found no significant difference in noise level across four face conditions (Experiment 1: *F*_*(3,45)*_ = 0.33, *p* = 0.80; Experiment 2-2: *F*_*(3,45)*_ = 0.23, *p* = 0.88).

We did the following statistical analyses: (1) we ran parametric repeated-measures ANOVAs with hemisphere (left and right OT ROIs), face condition (age, emotion, gender, and race), and SF/TF steps as factors on the categorical response amplitudes, with the JASP 0.95 software package. We were especially interested in measuring the hemispherical response differences in the categorization responses. (2) We then ran non-parametric bootstrap analyses to measure the minimal informational input threshold required for successful face categorization at different conditions. The details are as follows: a. 16 data points (i.e., 16 participants) were randomly drawn with replacement from each step and the mean of the drawn points was computed; b. the procedure was repeated 10,000 times; c. the mean and 95% confidence interval of the resulting distribution from step b. were calculated. The recognition threshold was defined as the first step at which the confidence interval exceeded zero.

#### EEG Time Domain Analysis

The spatio-temporal dynamics of the face categorization responses were investigated with similar methods as previous studies using the sweep SSVEP method ^14,15^. Referenced EEG signal was low-pass filtered with a 30 Hz cut-off (4^th^ order Butterworth filter), and cropped into an integer number of cycles of target face presentation frequency. After that, the general face presentation frequency and its harmonics (up to 30 Hz) were removed with narrow band notch-filtering (width = 0.1, slope = 0.1). The EEG waveforms were then segmented into smaller 1-s duration epochs with only one presentation of a target face. Epochs were then averaged and corrected relative to the baseline image(s) (200 ms) before the target face. This analysis was performed for each condition on individual participant before averaging into the group level.

To assess the statistical significance of neural responses relative to the baseline period (−200 to 0 ms), we defined a larger posterior region of interest (ROI) encompassing 19 electrodes over bilateral occipitotemporal and occipital cortex. This allowed us to test whether the frequency-domain findings generalized to a broader brain area ^13^. We performed a non-parametric cluster-based permutation test (10,000 iterations) on baseline-corrected data. First, single-sample t-tests (α = 0.05, two-tailed) identified significant time points (|*t*| > critical threshold corresponding to *p* < .05), which were then clustered based on temporal contiguity. Cluster strength was quantified as the cumulative sum of suprathreshold |t| values (cluster mass). By randomly permuting response polarities across participants, we generated a null distribution of maximal cluster masses, with observed clusters deemed significant when exceeding the 95^th^ percentile of this distribution (corrected *p* < 0.05) ^13^.

#### EEG time domain representational similarity analysis (RSA)

So far, we have examined mean categorization responses in each ROI at the group-level. Next, we examined the finer-grain representation of each face attribute within each individual’s brain by examining the spatiotemporal response patterns to each face condition. We adopted a time-resolved multivariate approach ^13^ and tested if neural representations in an individual’s brain were reliable across different trials of a condition using split-half analysis of the data, and if representations between conditions became more distinct as a function of increasing image spatial frequency content (Experiment 1) or image temporal exposure (Experiment 2). We predicted that if representations became more similar across trials within a condition and more dissimilar between those across conditions, distinctiveness would increase when enough visual exposure was presented.

For each condition, we computed mean temporal response profiles by integrating (concatenating) neural activity patterns across 19 posterior electrodes (the same electrodes as those selected in the group-level time domain analysis) within four consecutive 200-ms time windows (0-800 ms post-stimulus). Representational Similarity Matrices (RSMs) were then generated through split-half correlation analyses, enabling simultaneous assessment of both within-condition response stability and between-condition discriminability at the individual participant level. Category distinctiveness was defined as the difference between the within-category similarity (i.e., the correlation coefficient on the diagonal) of spatiotemporal responses across two split-half trials and the average between-category similarity (i.e., off-diagonal correlations) of spatiotemporal responses across trials ^13,17^. Distinctiveness is higher when the within-category similarity is bigger than the between-category similarity. We computed category distinctiveness for each face condition and in each participant across four above-mentioned time windows. Even though the distinctiveness index was derived from comparing within-vs. between-category correlations across blocks, our frequency-domain analysis of noise levels suggested that this measure was unlikely to be confounded by block-specific noise or SNR differences.

## Results

### Experiment 1

#### Categorical response amplitudes (1Hz and harmonics) as a function of increasing image spatial frequency content

In Experiment 1, we examined how measures of categorical response in four face attributes evolved as a function of increasing image spatial frequency content over nine incremental steps.

Figure 2. shows the averaged frequency spectra of face categorization responses (in signal-to-noise ratio, SNR) at 1 Hz (and its first 8 harmonics up to 9 Hz) averaged across four face conditions at each SF step over the OT ROI. Significant harmonics (above EEG noise) emerged at step 3, when images were low-pass filtered at 5.98 cycles/image (cpi). The recognition response mainly located over the posterior bilateral OT region, especially over the right hemisphere.

To investigate the categorization responses as a function of SF steps at face different conditions, we ran a three-way repeated measures ANOVA on the mean amplitude over the OT ROI with Hemisphere (left and right), Condition (age, emotion, gender, and race), and SF content (from step1 to step9) as within-subjects factors. The results showed significant main effects of Hemisphere, *F*_*(1, 15)*_ = 30.17, *p* < .001, *partial η*^*2*^ = .668, due to a right hemisphere advantage (left: M = .15 ± .28 μV, right: M = .27 ± .36 μV), and Condition, *F*_*(3, 45)*_ = 4.4, *p* = .024, *η*^*2*^ = .228, with a bigger response amplitude to race than gender (*t*_*(15)*_ = 4.595, *p* = .002, Bonferroni corrected; race: M = .28 ± .36; gender: M = .10 ± .25), and a bigger response amplitude to age than gender (*t*_*(15)*_ = 3.176, *p* = .038, Bonferroni corrected; age: M = .25 ± .35; gender: M = .10 ± .25), and SF content, *F*_*(8, 120)*_ = 18.56, *p* < .001, *η*^*2*^ = .553. There were also significant two-way interactions of Hemisphere × Condition, *F*_*(3, 45)*_ = 4.87, *p* = .005, *η*^*2*^ = .245, Hemisphere × SF content, *F*_*(8, 120)*_ = 5.60, *p* < .001, *η*^*2*^ = .27, and Condition × SF content, *F*_*(24, 360)*_ = 2.15, *p* = .002, *η*^*2*^ = .125. The three-way interaction of Hemisphere × Condition× SF content was not significant, *F*_*(24*, 360)_ = 1.45, *p* = .080, *η*^*2*^ = .088.

Next, we ran non-parametric bootstrap analysis separately for each face condition to find out the minimal amount of SF content (i.e. starting to be significantly above zero) capable of driving successful face categorization (see Methods part). The response amplitudes were measured in the right OT (rOT) ROI only, as the ANOVA revealed a right hemisphere advantage in all face categorization tasks. Among the four conditions, significant responses emerged when images were filtered at 4.89 cpi for race and emotion, followed by age at 7.31 cpi (Figure 3A). Gender categorization emerged the latest when images were filtered at 10.94 cpi. Scalp topographies (Figure 3B) revealed robust activation patterns in the lateral occipitotemporal (OT) electrodes, which systematically increased with image content strength. Notably, these activation patterns demonstrated consistency across all experimental conditions, suggesting that common neural populations in the occipitotemporal cortex were engaged for face attributes processing – independent of spatial frequency variations.

**Figure 3.**
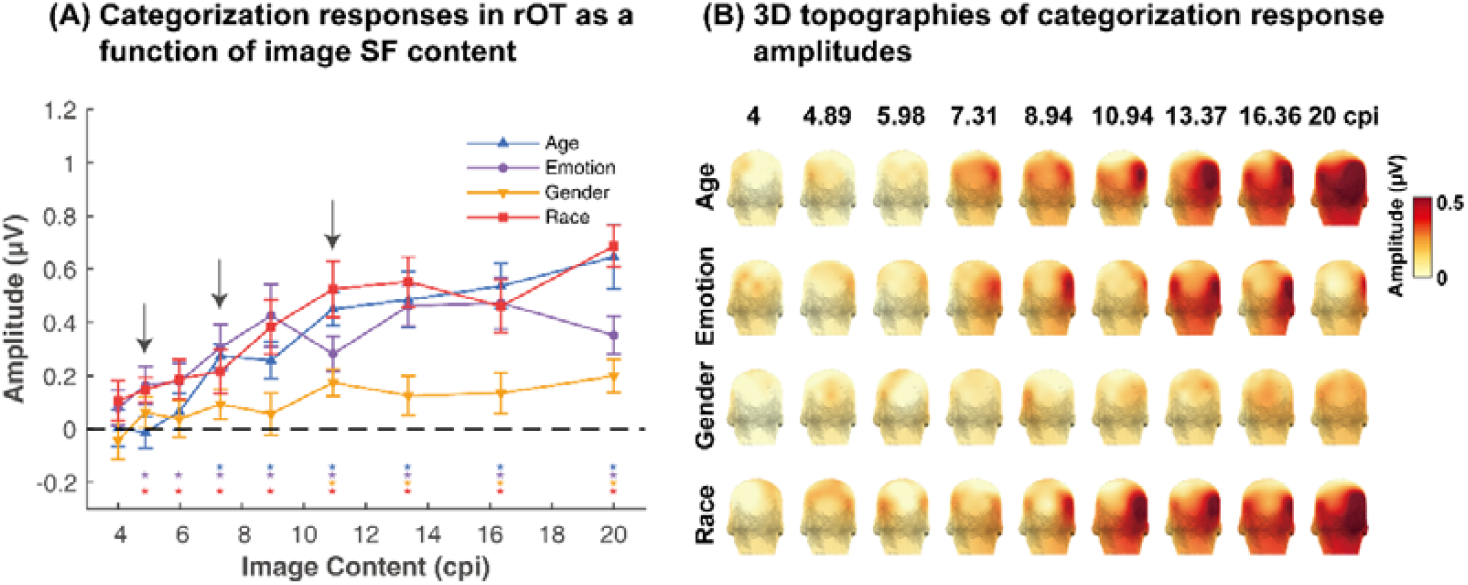
Face categorization response profiles as a function of image SF content for all conditions. **(A)** Response amplitudes in the rOT ROI. Errors: 95% confidence interval. Asterisks: significant categorization responses from zero. Black arrows: the emergence of significant face categorization responses. **(B)** Three-D topographies.

**Temporal dynamics reveal different spatiotemporal response patterns across conditions Figure 4** presents the time domain results over 19 posterior electrodes across four face conditions. Time series are averaged across several steps according to the frequency-domain findings (**Figure 3A**): steps 2-3 (4.89-5.98 cpi) as race and emotion categorization emerged in these two steps, steps 4-5 (7.31-8.94 cpi) as age categorization emerged in these two steps, and steps 6-9 (10.94-20 cpi) where categorization responses in all conditions were significant. Our analyses revealed three key findings: (1) While spatiotemporal dynamics varied between conditions, they remained remarkably consistent within each condition across SF steps. (2) Significant categorization responses consistently occurred 200-600 ms post-stimulus across all conditions. (3) The condition-specific response patterns suggested differential neural processing dynamics for distinct facial attributes, despite shared engagement of posterior brain regions.

**Figure 4.**
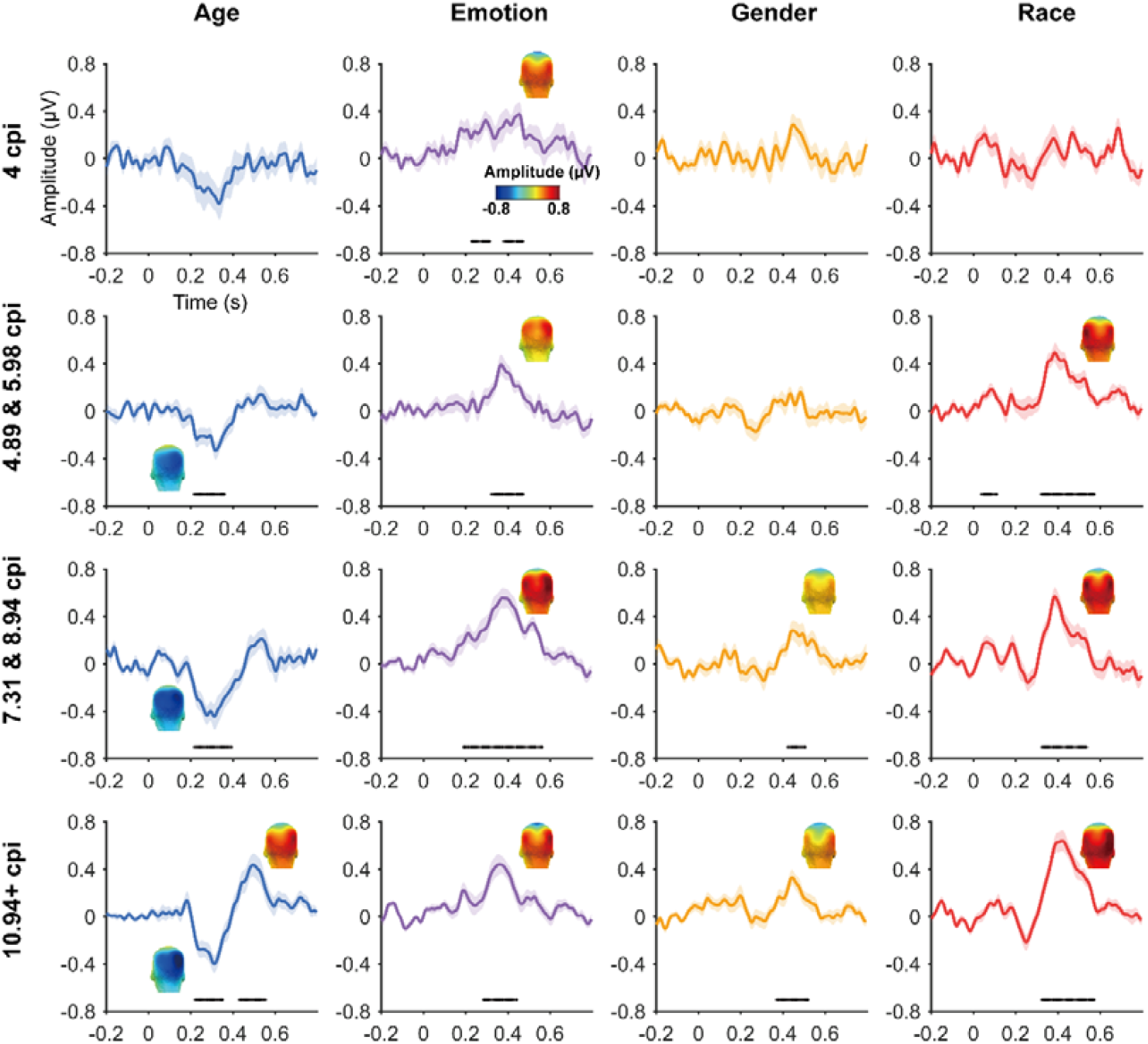
Temporal dynamics of category-selective responses as a function of image SF content. Data are averaged across electrodes of a larger posterior ROI across individuals and different SF steps. Colored lines: mean responses. Shadings: standard errors across participants. Colored horizontal lines at y = ×0.7: significant responses relative to zero for the posterior ROI (*ps* < .05). 3D topographies: spatial distribution of the face categorization responses at significant time windows [peak latency ± 10 ms].

#### Representational similarity analyses reveal different modulation effect of image spatial frequency content on face attribute perception

Our observation of distinct spatiotemporal dynamics over posterior brain regions for the four face conditions (**Figure 4**) prompted an investigation into the trial-level reliability of fine-grained category representations within individual participant. We predicted that progressive clarity of facial features would enhance representation similarity between odd/even split-half trials within each category condition, but increase dissimilarity between different categories.

To examine the representational structure, we calculated representation similarity matrices (RSMs) across odd/even split-halves of the data in each participant and each condition (**Figure 5**). Each cell in the RSM quantified the similarity between two spatiotemporal patterns: On-diagonal cells of RSM quantify the similarity of distributed spatiotemporal responses to odd/even trials from the same category condition and off-diagonal cells quantify the similarity of spatiotemporal responses to odd/even trials from different categories.

**Figure 5.**
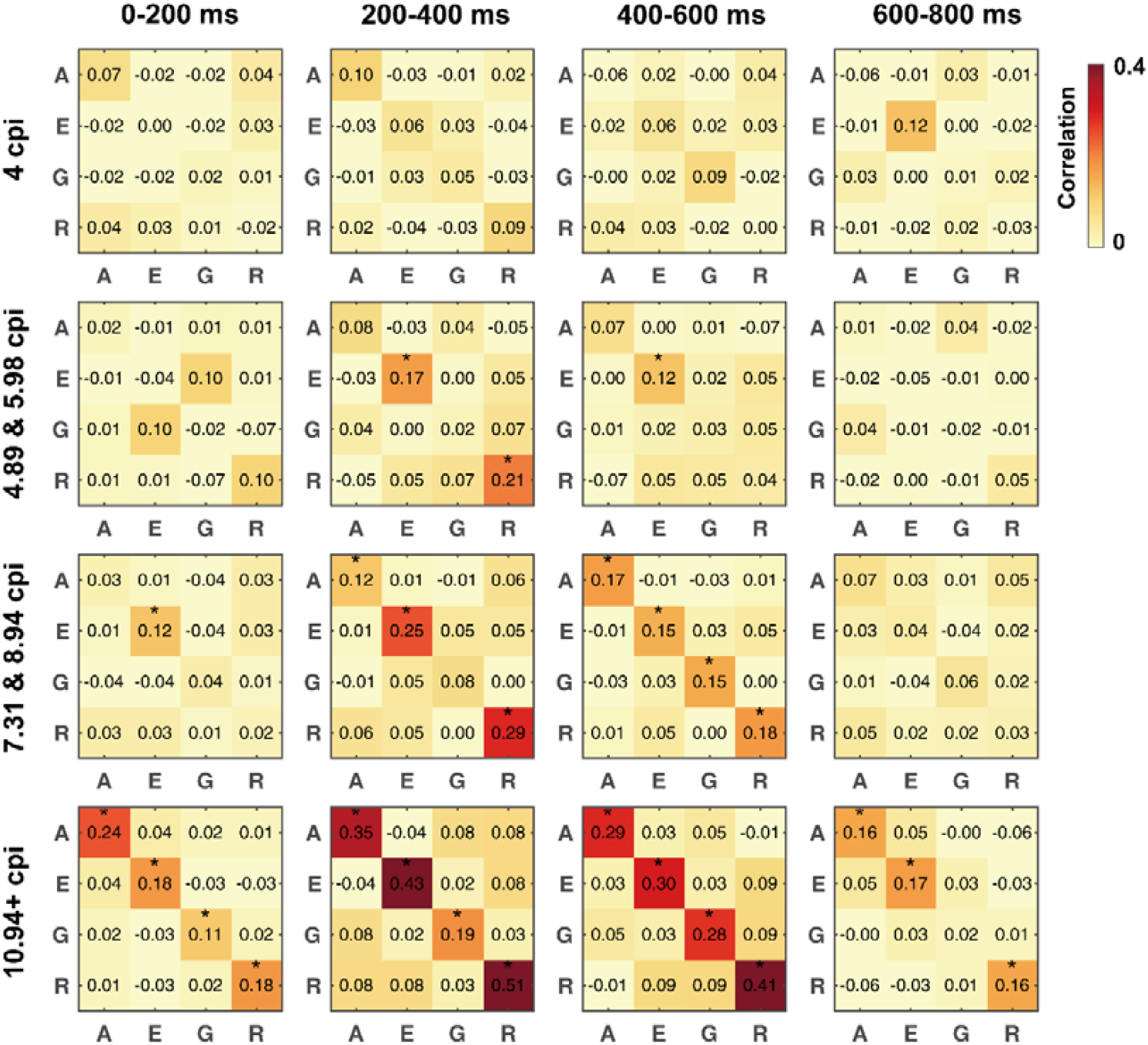
Representation similarity matrices (RSMs) generated from odd/even split-halves of the spatiotemporal patterns of categorization responses. Spatiotemporal patterns for each category were generated by concatenating the mean time courses of 19 electrodes covering the posterior brain region. Asterisks: significant above-zero within-condition correlations between split-half trials (*ps* < .05, one-tailed; FDR-corrected at 4 levels). A: age. E: emotion. G: gender. R: race.

Examination of mean RSM in each condition revealed that within-category similarity was not significantly above zero with very blurry faces (4 cpi) across four time windows (**Supplementary Table S1, Figure 5**). When SF content of images increased to 4.89 and 5.98 cpi, distributed responses to emotion and race became reliable, as within-category similarity for these two categories was significantly above zero in 200-400 ms (*ps* < .01, one-tailed; FDR corrected). A significant within-category similarity was also found for emotion in 400-600 ms (*p* = .04). When SF content increased to 7.31 and 8.94 cpi, emotion was significant in 0-200 ms (*p* = .02). Age, emotion and race were significant in 200-400 ms (*ps* < .05). All conditions were significant in 400-600 ms (*ps* < .05). When SF content increased to 10.94 cpi and above, distributed responses to all conditions became reliable in all time windows (*ps* < .05), except gender in the last time window.

Next, we evaluated the category distinctiveness as a function of image SF content. When faces were very blurry (4cpi), distinctiveness score in all conditions were not significantly above zero, suggesting no differences between spatiotemporal responses to one category vs. another (**Supplementary Table S2, Figure 6**). When SF content increased to 4.89 – 5.98 cpi, category distinctiveness appeared in emotion and race condition in 200-400 ms (*ps* < .05). When SF content increased to 7.31 – 8.94 cpi, emotion and race were significant in 200-400 ms (*ps* < .01). All conditions were significant in 400-600 ms (*ps* < .05). When SF content increased to 10.94 cpi and above, all conditions became significant (*ps* < .05) in all time windows except gender in 600-800 ms.

**Figure 6.**
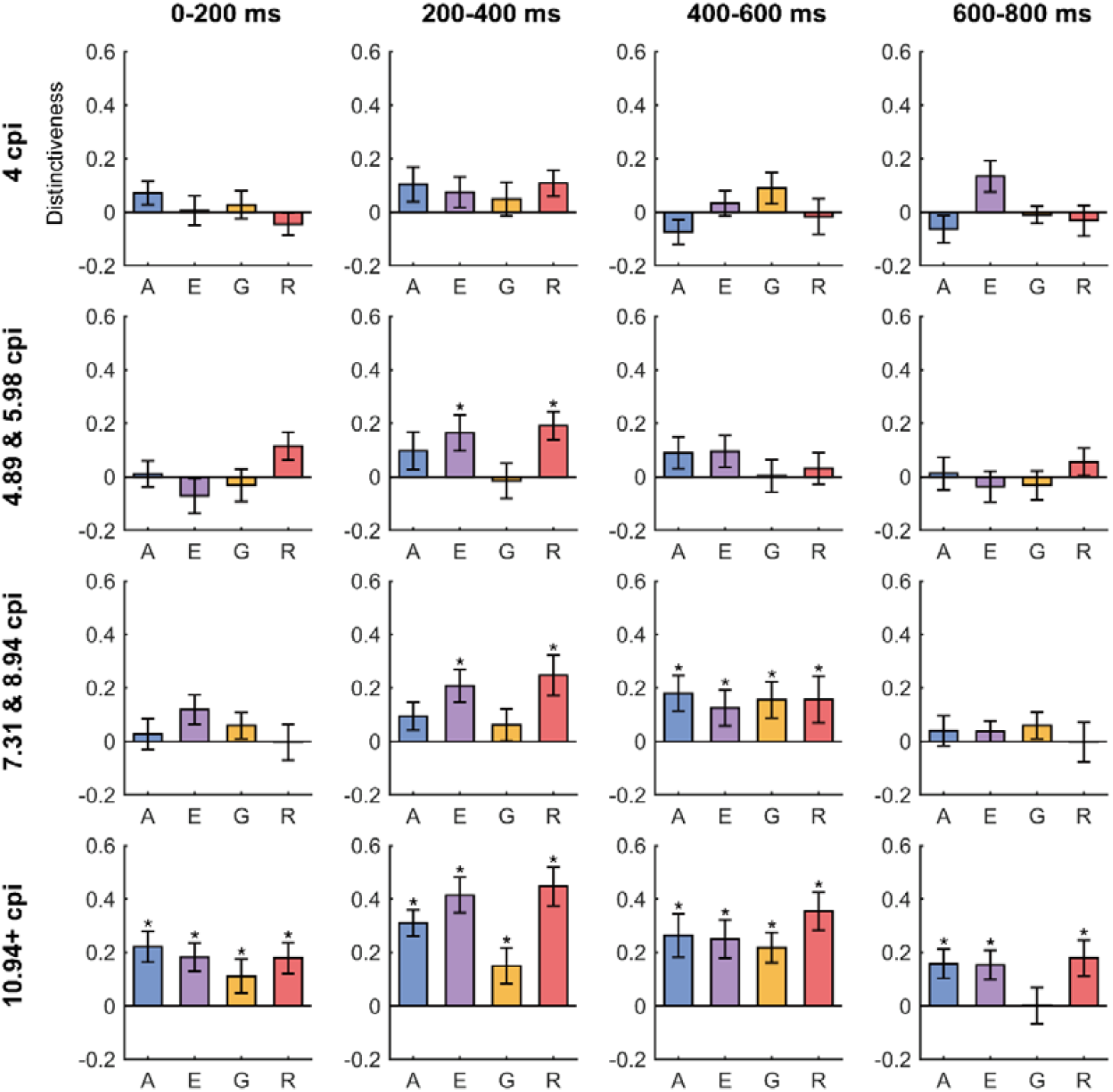
Category distinctiveness scores calculated for each participant and face condition by subtracting the mean between-category correlation values from the within-category correlation values between split-half trials. Asterisks: significant above-zero distinctiveness scores (*ps* < .05, FDR corrected at 4 levels). A: age. E: emotion. G: gender. R: race.

## Experiment 2

### Categorical response amplitudes (1 Hz and harmonics) as a function of increasing image exposure time

We ran a preliminary experiment from 10 participants to examine face categorization responses in four conditions when images were exposed from 12 Hz to 3 Hz in seven steps. The results showed significant categorization responses in all four face conditions even when faces were presented for only 83 ms each (12 Hz; full results are documented in the supplementary file), therefore, in the follow-up Experiment 2-2, we increased the maximum image presentation rate to 24 Hz, and measured how face categorization responses evolved as a function of increasing image exposure time (i.e., from 24 Hz to 12 Hz over four incremental steps).

Similar to Experiment 1, significant categorization responses were found at 1 Hz and harmonics over the OT ROI, especially when images were presented at a slower rate (**Supplementary Figure 3**). To investigate the face categorization responses as a function of TF steps at different conditions, we ran a three-way repeated-measures ANOVA on the mean amplitude over the OT ROI with Hemisphere (left, and right), Condition (age, emotion, gender, and race), and TF content (4 steps) as within-subjects factors. The results showed significant main effects of Hemisphere, *F*_*(1, 15)*_ = 6.25, *p* = .024, *partial η*^*2*^ = .294, due to a right hemisphere advantage (left: M = .10 ± .21 μV ; right M = .15 ± 0.25 μV), and TF content, *F*_*(3, 45)*_ = 13.51, *p* < .001, *η*^*2*^ = .474. There was also a significant two-way interaction of Condition × TF content, *F*_*(9, 135)*_ = 2.01, *p* = .043, *η*^*2*^ = .118. The three-way interaction of Hemisphere × Condition × TF content was not significant, *F*_*(9, 135)*_ = 1.22, *p* = .289, *η*^*2*^ = .075. No further effects were significant (all *ps* > .1).

We then ran a non-parametric bootstrap analysis separately for each condition in the right OT ROI to find out the minimal amount of image duration (i.e. starting to be significantly above zero) capable of driving successful face categorization at different attributes (see Methods part). Significant categorization responses were observed when images were presented at 41.67 ms for age, and emotion, followed by a significant response to race at 50 ms exposure and gender later at 67 ms (**Figure 7A**). Inspection of the 3-D scalp topographies (**Figure 7B**) showed activations of lateral occipitotemporal regions especially in the right hemisphere across increasing image duration.

**Figure 7.**
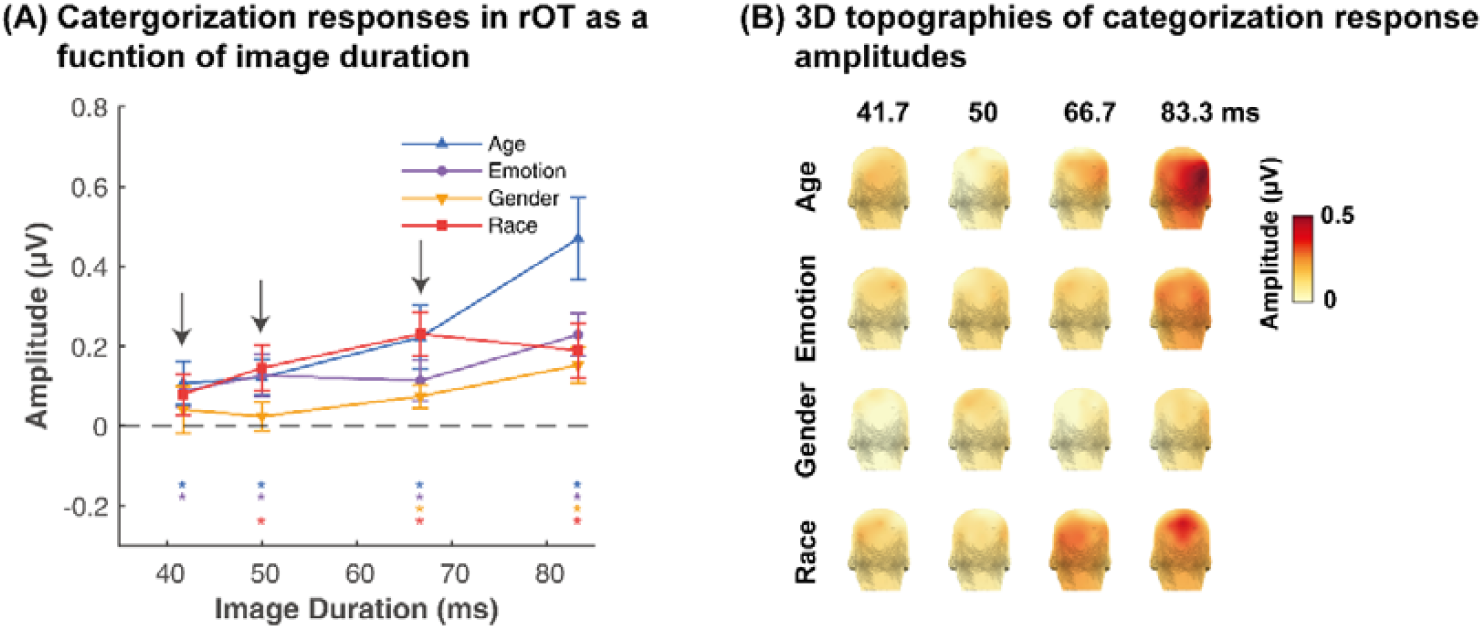
Face categorization response profiles as a function of image duration for all face conditions. **(A)**. Response amplitudes in the rOT ROI. Errors: 95% confidence interval. Asterisks: significant categorization responses from zero (*ps* < .05, FDR-corrected). Black arrows: the emergence of significant categorization responses. **(B)**. Three-D topographies.

### Representational similarity analyses reveal different modulation effect of image duration on face attribute perception

Time-domain analyses confirmed distinct spatiotemporal dynamics across face conditions (**Supplementary Figure 4**). To systematically compare these response patterns, we quantified representational similarity of distributed neural responses using split-half trials, assessing both within- and between-condition similarities. Examination of mean RSMs in each face condition showed that within-condition similarity for race was significantly above zero in 0-200 ms (*p* = .011, one-tailed; FDR-corrected) and age in 400-600 ms (*p* = .04) when images were presented at 41.7 ms (**Supplementary Table S3, Figure 8A**). When image duration increased to 50 ms, race was significant in 0-200 ms (*p* = .004), emotion and race were significant in 200-400 ms (*ps* < .05). When image duration increased to 66.7 - 83.3 ms, distributed responses to all face conditions became reliable in 0-400 ms (*ps* < .001), except gender in 200-400 ms.

**Figure 8.**
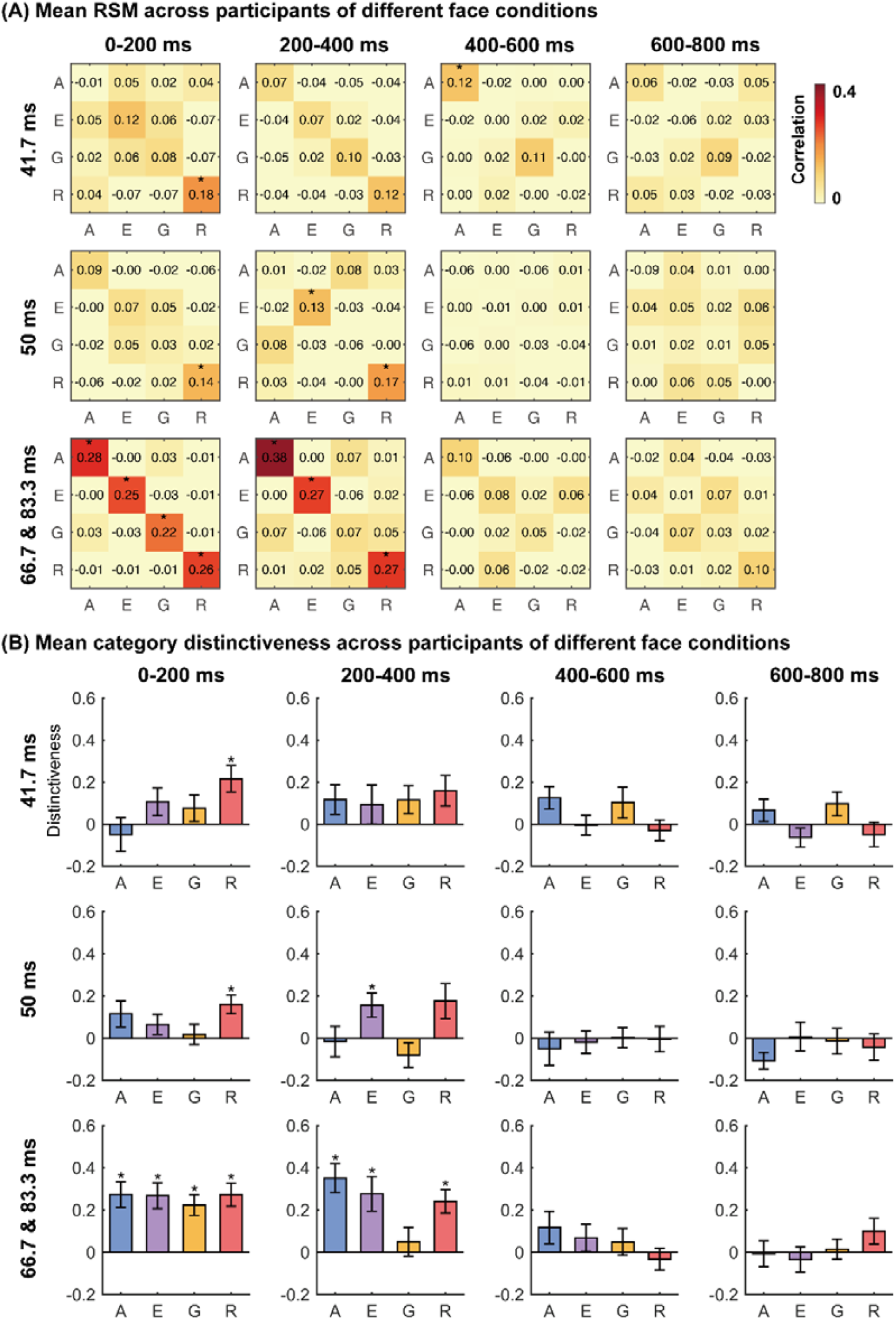
Representation similarity matrices (RSMs) generated from odd/even split-halves of the spatiotemporal patterns of categorization responses. **(A)**. RSMs generated from odd/even split-halves of the spatiotemporal patterns of responses. Spatiotemporal patterns for each category were generated by concatenating the mean time-courses of 19 electrodes covering the posterior brain region. Asterisks: significant above-zero within-condition correlations between split-half trials (*ps* < .05; FDR-corrected at 4 levels). **(B)**. Category distinctiveness scores calculated for each participant and face condition by subtracting the mean between-category correlation values from the within-category correlation values between split-half trials. Asterisks: significant above-zero distinctiveness scores (*ps* < .05, FDR corrected at 4 levels). A: age. E: emotion. G: gender. R: race.

Furthermore, category distinctiveness appeared in race in 0-200 ms (*p* = .007; FDR-corrected) when images were presented at 41.7 ms (**Supplementary Table S4, Figure 8B**). When image duration increased to 50 ms, race was significant in 0-200 ms (*p* = .005) and emotion in 200-400 ms (*p* = .031). When image duration increased to 66.7 ms and above, category distinctiveness scores in all conditions were significant (*ps* < .01), except gender in 200-400 ms.

## Discussion

This study investigates how the brain dynamically processes face attributes (age, emotion, gender, and race) under two ecologically relevant scenarios: (1) faces gradually sharpening from blur to clear (mimicking approaching from a distance), and (2) faces presented with incrementally increasing temporal exposure (mimicking brief, flashed encounters). Our results reveal that emotion and race recognition require less spatial or temporal exposure, whereas age categorization shows context-dependent processing: it demands more SF details under low-spatial-detail conditions but requires less exposure time when facial details are available. Representational similarity analyses further show that reliable representations of face attributes are modulated differently by the visual constraints. Together, these results reveal a flexible **temporal processing system** in which the brain adaptively prioritizes different facial features based on available visual input, offering novel insights into how naturalistic visual challenges are resolved through dynamic neurocomputational mechanisms.

To our knowledge, this study provides the first systematic investigation of flexible perception of face attributes under constrained visual conditions that mimic daily scenarios. Previous studies have attempted to establish a temporal processing order (e.g., age → emotion → identity in ^5^) under optimal viewing conditions (e.g., using full-spectrum images with exposures of ≥ 200 ms), but they presented isolated, unmasked faces that lack ecological validity and critically omitted race processing ^3,5^. By presenting clear images with much shorter temporal durations, our results offer three key advances: (1) remarkably efficient extraction of age and race information, with reliable above-chance categorization at just 41.67 ms exposure; (2) relatively rapid emotion processing emerging by 50 ms exposure; and (3) confirmation of a canonical age/race → emotion temporal processing order that was partly consistent with ^5^ ‘s framework. Although face identity was not examined in the present study, extensive literature supports its later processing timeline ^2,3,5,18^, as well as its modulation by other facial attributes ^2,19^. The efficient processing of age information observed here is also consistent with previous findings ^3,20^.

However, when face images were presented with sufficient temporal exposure (e.g., 200 ms) but under spatial content constrained conditions, results from Experiment 1 revealed distinct temporal order of face processing: (1) emotion and race information were efficiently extracted at very low spatial frequencies (∼5.98 cpi), likely mediated by coarse structural cues; (2) age categorization, by contrast, was delayed (∼7.31 cpi) due to its reliance on finer facial details. Notably, despite this delay, age processing still precedes identity recognition (which requires 8-16 cpi SF resolution according to previous findings, e.g., ^10^ with the same sweep SSVEP paradigm, and ^21–33^); (3) confirmation of a novel computational Emotion/Race → Age temporal order. Taken together, results from both experiments demonstrate how the brain dynamically prioritizes different facial attributes based on available visual input, highlighting the need to refine existing face perception models.

To investigate the reliable representation of facial attributes within individual participants, we applied time-domain Representational Similarity Analysis (RSA) across an extended posterior brain region ^13,17^. This analysis revealed that the temporal order in which reliable representations emerged for face attributes under different visual constraint conditions were not fixed (Figure 5,6,8). This key finding, demonstrating a flexible processing sequence, was consistent with the results of the ROI-based group analysis, thereby substantiating the effect at both the individual and group levels. Under conditions of constrained spatial content, reliable spatiotemporal response patterns for emotion and race emerged early, with within-condition split-half correlations significantly exceeding between-condition correlations from 200–600 ms post-stimulus. In contrast, reliable representations for age and gender emerged later, indicating their greater dependence on finer spatial frequency details for robust neural coding. When images were low-pass filtered at ≥ 10.95 cpi, all face attributes (except gender in 600–800 ms window) exhibited distinct spatiotemporal patterns across all time windows. Under temporal constraints, reliable age and race representations emerged the earliest (by 41.67 ms exposure), followed by emotion (50 ms) and gender (≥ 66.7 ms). Notably, significant decoding windows under temporal constraints appeared earlier (i.e., 0-400 ms in Experiment 2) than those under spatial content constraints (i.e., 200-600 ms in Experiment 1), suggesting a more rapid onset of information processing relative to stimulus onset. This temporal pattern was further supported by data from Experiment 2-1, which showed that the peak latency of a negative deflection in the race condition, as well as a positive deflection in the emotion condition, increased systematically with longer image durations (**Supplementary Figure S3**).

Our study, consistent with findings from ^5^, observed weaker neural responses to gender information. In fact, decoding performance in ^5^ even fell below chance level, suggesting that gender processing (at least for Asian faces) heavily relies on external cues (e.g., hairstyle, makeup, facial hair). Supporting this, ^3^ reported an early detection of gender information when such cues were present. When these external cues were removed, our results showed that gender categorization required more sensory input (i.e., more spatial details in Experiment 1, and longer temporal exposure in Experiment 2) than the other three face attributes. Results of our study contrast with earlier behavioral studies that reported near-ceiling gender categorization accuracy even with minimized external cues under prolonged viewing durations (e.g., ≥ 1000 ms) using photographs or 3D laser-scanned faces ^34–36^. Those studies argue that features such as eyebrows and skin texture play an important role in gender categorization. The discrepancy between our findings and prior behavioral results suggests that different experimental paradigms may engage distinct computational strategies in gender processing. An interesting direction for future research will be to investigate whether factors like face race or age are interacted with gender categorization.

What are the important features that drive early and reliable categorization on age, emotion, and race information from a face? Results of our two experiments indicate that coarse shape information plays a key role in emotion and race categorization, especially when facial textures are unavailable under blurry conditions. Instead for age categorization, details on the faces, such as skin texture or wrinkles, are more important ^37^. To mechanistically identify the critical features underlying efficient face categorization, future work could employ deep convolutional neural networks (DCNNs). These models are well-suited for this purpose, as they achieve near-human performance in face recognition and replicate the hierarchical processing patterns of the primate visual system ^38,39^. A powerful approach would be to correlate the representational dissimilarity matrix (RDM) from each EEG time point with the RDM from each progressive layer of a DCNN processing the same stimuli ^3,7,40^. At time points where neural and model representations are significantly aligned, feature visualization techniques ^41–43^ can be applied to the corresponding DCNN layers. This would enable a precise mapping of how specific convolutional kernels encode diagnostic features, such as emotion-relevant contours or age-predictive textures, thereby revealing the computational strategies that underlie rapid human categorization.

Besides the temporal dynamics of face perception, understanding the spatial organization of face processing remains an important question in human neuroscience. Functional MRI studies reveal dissociable pathways for processing facial emotion (amygdala and posterior superior temporal sulcus, pSTS) versus identity (fusiform face area, FFA, and anterior inferior temporal cortex, aIT), with preferential decoding of emotion in amygdala and pSTS and identity in the ventral FFA and aIT brain region ^44–48^. Other face attributes including gender and age are also encoded in the right-lateralized occipital and fusiform face areas (rOFA/rFFA) ^49–51^. Future studies combining intracranial EEG ^4^ or simultaneous EEG/MEG-fMRI ^52–56^ could further elucidate both the millisecond-scale dynamics and brain activation networks underlying this flexible temporal processing schedule.

While our study advances the understanding of flexible, temporal processing of face attributes, a potential limit lies in the exclusive use of young Asian and Caucasian female faces for examining race categorization (a constraint that also applies to other facial attributes due to the use of limited range of exemplars). Although this design choice helped minimize total experimental time but at the same time offered high SNR, it introduces the possibility that race-related cues (or cues from other face attributes) may be more salient in young female faces than in other demographic groups (e.g., old or male faces), hence biases the results. However, the consistency between our findings in Experiment 2-2 and those of prior work ^5^ (using fully balanced stimuli) suggests that such bias may be minimal.

Similarly, the early emergence of emotion categorization we observed may be influenced by our stimulus set, which contrasted only happy and sad faces. The pronounced difference in mouth aperture (e.g., open mouths/teeth in happy faces versus closed mouths in sad faces) provides a salient visual cue that could facilitate rapid emotion detection. However, the response advantage for one emotion over another is often subtle and context-dependent. Previous EEG studies have reported small response differences between some facial expressions but not all, such as larger amplitudes for sadness compared to fear or happiness ^57^, or for disgust compared to happiness ^58^. Furthermore, decoding onset latencies are typically confined to an 80-120 ms window ^7,59,60^, although significant differences have also been reported. For instance, happy and fearful faces can be decoded significantly earlier (< 100 ms) than angry faces (at approximately 220 ms) ^52^. When examining the minimal temporal exposure for facial expression processing, a recent study did not find any emotion modulation effect, either ^61^. Future studies could use randomized stimulus sets in which non-target features (e.g., age, gender, emotion) are varied, while the target category (e.g., race) is systematically controlled between target and baseline images. This approach would help mitigate potential confounds in face categorization while maintaining ecological relevance.

In summary, by combining sweep SSVEP paradigms, we investigated how perception of face attributes (e.g., age, emotion, gender, and race) dynamically adapts to ecologically relevant scenarios. Our results revealed a flexible, temporal processing system in which the brain selectively prioritizes different facial attributes based on available visual input. These findings advance our understanding of visual processing by highlighting adaptive neurocomputational mechanisms and offer a unified framework for studying dynamic face perception under naturalistic visual constraints.

## Data availability statement

Individual preprocessed EEG data, behavioral data, and code for all analyses are available on OSF (https://osf.io/w7efy/).

## Acknowledgements

This work was supported by grants from the National Natural Science Foundation of China (NSFC) (No. 32300860) to X.Y.

## Authorship contributions

S.L., Data collection, Formal analysis, Validation, Visualization, Methodology, Writing; Y.W., Data collection; X.Y., Conceptualization, Resources, Formal analysis, Software, Supervision, Funding acquisition, Methodology, Writing.

