## Supplementary for "Flexible perception of face attributes under naturalistic visual constraints"

**Supplementary information**

**1. Image analysis**

In the experiments, we combined the sweep SSVEP paradigm to measure the minimal sensory input required for implicit face attributes categorization. All visual stimuli were normalized for luminance and contrast to mitigate the influence of low-level image cues on high-level facial processing. No further image manipulation was applied. However, we further ran spatial frequency analysis on the images and showed that the images possessed similar spectral power distributions across categories (**Figure S1**).

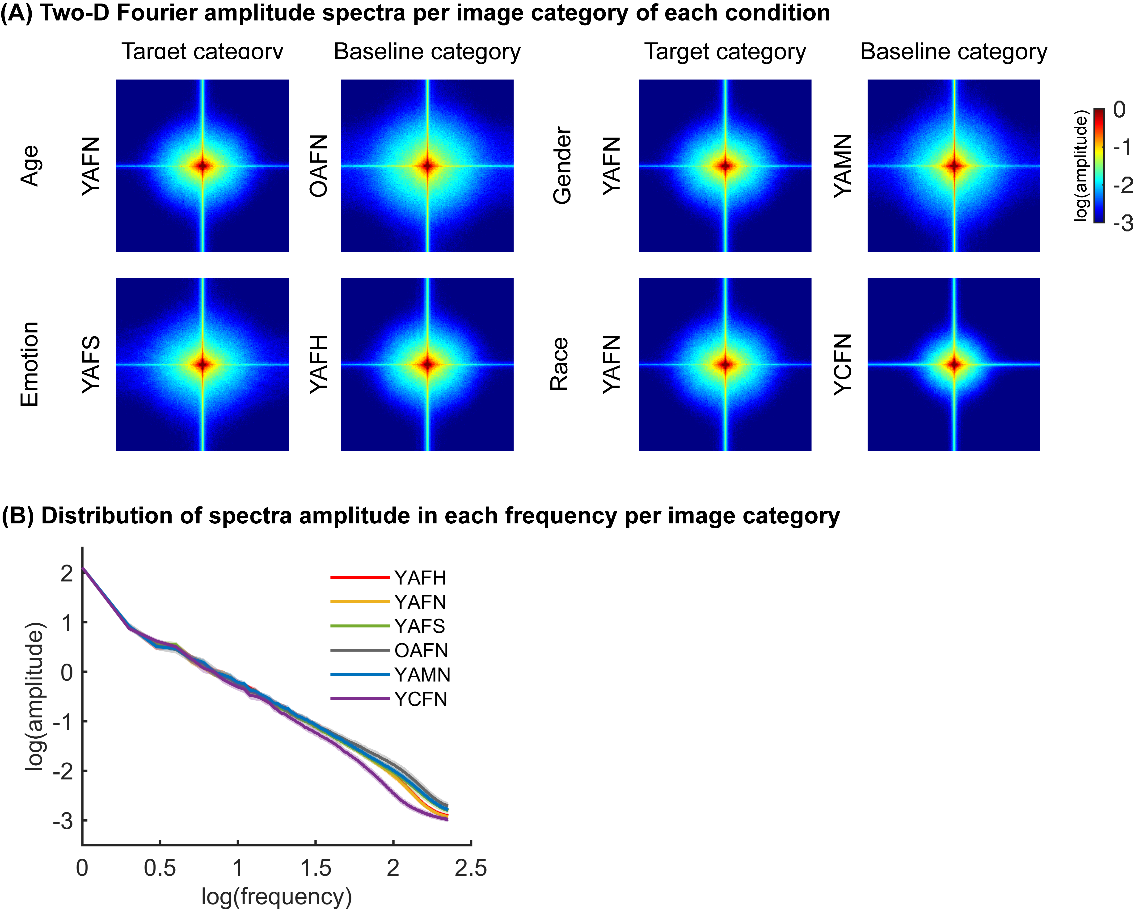

**Figure S1. Image spatial frequency analysis.** (A) Two-D Fourier amplitude spectra (log transformed) averaged across 21 images in each category. (B) Distribution of spectral amplitude in each frequency averaged across 21 images in each category. Shaded areas: standard deviation. H: Happy, Y: Young, A: Asian, F: Female, N: Neutral, S: Sad, M: Male, O: Old, C: Caucasian.

**2. Experiment 1: spatial frequency sweep SSVEP task**

**Table S1. Statistical results for on-diagonal within-condition correlations between odd/even trials in the RSMs across four time windows.** Bold texts: significant above-zero within-condition correlations (FDR-corrected).

| Age | 0-200 ms | 200-400 ms | 400-600 ms | 600-800 ms |
| --- | --- | --- | --- | --- |
| 4 cpi | *t_(15)_* = 1.99, *p* = .129,  *d* = 0.498 | *t_(15)_* = 2.09, *p* = .071,  *d* = 0.522 | *t_(15)_* = -1.31, *p* = .896,  *d* = -0.328 | *t_(15)_* = -1.40, *p* = .909,  *d* = -0.350 |
| 4.89 & 5.98 cpi | *t_(15)_* = 0.39, *p* = .699,  *d* = 0.099 | *t_(15)_* = 1.41, *p* = .120,  *d* = 0.352 | *t_(15)_* = 1.29, *p* = .218,  *d* = 0.322 | *t_(15)_* = 0.23, *p* = .811,  *d* = 0.056 |
| 7.31 & 8.94 cpi | *t_(15)_* = 0.59, *p* = .352  *d* = 0.147 | ***t_(15)_* = 2.69, *p =* .011,**  ***d* = 0.672** | ***t_(15)_* = 2.92, *p =* .012,**  ***d* = 0.731** | *t_(15)_* = 1.45, *p* = .190,  *d* = 0.363 |
| 10.94 + cpi | ***t_(15)_* = 4.69, *p* = 4.8*10^-4^,**  ***d* = 1.173** | ***t_(15)_* = 10.31, *p* = 8*10^-8^,**  ***d* = 2.577** | ***t_(15)_* = 3.89, *p* = 7.2*10^-4^,**  ***d* = 0.973** | ***t_(15)_* = 3.37, *p* = .004,**  ***d* = 0.843** |
| Emotion | 0-200 ms | 200-400 ms | 400-600 ms | 600-800 ms |
| 4 cpi | *t_(15)_* = 0.02, *p* = .657,  *d* = 0.005 | *t_(15)_* = 1.46, *p* = .109,  *d* = 0.366 | *t_(15)_* = 1.35, *p* = .198,  *d* = 0.337 | *t_(15)_* = 2.24, *p* = .081,  *d* = 0.561 |
| 4.89 & 5.98 cpi | *t_(15)_* = -0.79, *p* = .780,  *d* = -0.198 | ***t_(15)_* = 2.78, *p* = .014,**  ***d* = 0.696** | ***t_(15)_* = 2.60, *p* = .040,**  ***d* = 0.650** | *t_(15)_* = -0.91, *p* = .811,  *d* = -0.227 |
| 7.31 & 8.94 cpi | ***t_(15)_* = 2.96, *p* = .020,**  *d* = 0.739 | ***t_(15)_* = 4.60, *p* = 3.5*10^-4^,**  ***d* = 1.149** | ***t_(15)_* = 2.62, *p* = .012,**  ***d* = 0.655** | *t_(15)_* = 1.11, *p* = .190,  *d* = 0.277 |
| 10.94 + cpi | ***t_(15)_* = 4.44, *p* = 4.8*10^-4^,**  ***d* = 1.110** | ***t_(15)_* = 8.91, *p* = 1.5*10^-7^,**  ***d* = 2.228** | ***t_(15)_* = 4.45, *p* = 3.1*10^-4^,**  ***d* = 1.112** | ***t_(15)_* = 3.84, *p* = .003 ,**  ***d* = 0.961** |
| Gender | 0-200 ms | 200-400 ms | 400-600 ms | 600-800 ms |
| 4 cpi | *t_(15)_* = 0.37, *p* = .657,  *d* = 0.092 | *t_(15)_* = 0.83, *p* = .209,  *d* = 0.208 | *t_(15)_* = 1.79, *p* = .187,  *d* = 0.448 | *t_(15)_* = 0.31, *p* = .764,  *d* = 0.076 |
| 4.89 & 5.98 cpi | *t_(15)_* = -0.37, *p* = .780,  *d* = -0.092 | *t_(15)_* = 0.38, *p* = .354,  *d* = 0.096 | *t_(15)_* = 0.61, *p* = .274,  *d* = 0.154 | *t_(15)_* = -0.53, *p* = .811,  *d* = -0.133 |
| 7.31 & 8.94 cpi | *t_(15)_* = 0.76, *p* = .352,  *d* = 0.190 | *t_(15)_* = 1.49, *p* = .078,  *d* = 0.373 | ***t_(15)_* = 2.87, *p* = .012,**  ***d* = 0.718** | *t_(15)_* = 1.33, *p* = .190,  *d* = 0.333 |
| 10.94 + cpi | ***t_(15)_* = 2.26, *p* = .020,**  ***d* = 0.566** | ***t_(15)_* = 3.26, *p* = .003,**  ***d* = 0.815** | ***t_(15)_* = 5.12, *p* = 1.3*10^-4,^**  ***d* = 1.281** | *t_(15)_* = 0.27, *p* = .395,  *d* = 0.068 |
| Race | 0-200 ms | 200-400 ms | 400-600 ms | 600-800 ms |
| 4 cpi | *t_(15)_* = -0.59, *p* = .717,  *d* = -0.147 | *t_(15)_* = 1.94, *p* = .071,  *d* = 0.486 | *t_(15)_* = 0.01, *p* = .661,  *d* = 0.003 | *t_(15)_* = -0.70, *p* = .909,  *d* = -0.175 |
| 4.89 & 5.98 cpi | *t_(15)_* = 2.40, *p* = .060,  *d* = 0.600 | ***t_(15)_* = 4.49, *p* = 8.6*10^-4^,**  ***d* = 1.124** | *t_(15)_* = 0.79, *p* = .274,  *d* = 0.198 | *t_(15)_* = 1.25, *p* = .461,  *d* = 0.312 |
| 7.31 & 8.94 cpi | *t_(15)_* = 0.39, *p* = .352,  *d* = 0.097 | ***t_(15)_* = 4.90, *p* = 3.5*10^-4^,**  ***d* = 1.224** | ***t_(15)_* = 2.50, *p* = .012,**  ***d* = 0.625** | *t_(15)_* = 0.42, *p* = .340,  0.105 |
| 10.94 + cpi | ***t_(15)_* = 3.85, *p* = .001,**  ***d* = 0.964** | ***t_(15)_* = 8.96, *p* = 1.5*10^-7^,**  ***d* = 2.239** | ***t_(15)_* = 7.30, *p* = 5.2*10^-6^,**  ***d* = 1.826** | ***t_(15)_* = 3.21, *p* = .004,**  ***d* = 0.802** |

**Table S2. Statistical results for distinctiveness scores for each face condition across four time windows in Experiment 1.** Bold texts: significant within- vs. between-category similarities (FDR-corrected).

| Age | 0-200 ms | 200-400 ms | 400-600 ms | 600-800 ms |
| --- | --- | --- | --- | --- |
| 4 cpi | *t_(15)_* = 1.67, *p* = .231,  *d* = 0.418 | *t_(15)_* = 1.61, *p* = .128,  *d* = 0.403 | *t_(15)_* = -1.60, *p* = .935,  *d* = -0.400 | *t_(15)_* = -1.23, *p* = .881,  *d* = -0.307 |
| 4.89 & 5.98 cpi | *t_(15)_* = 0.23, *p* = .821,  *d* = 0.057 | *t_(15)_* = 1.38, *p* = .125,  *d* = 0.345 | *t_(15)_* = 1.50, *p* = .154,  *d* = 0.376 | *t_(15)_* = 0.22, *p* = .732,  *d* = 0.055 |
| 7.31 & 8.94 cpi | *t_(15)_* = 0.48, *p* = .426,  *d* = 0.120 | *t_(15)_* = 1.80, *p* = .061,  *d* = 0.450 | ***t_(15)_* = 2.72, *p* = .032,**  ***d* = 0.680** | *t_(15)_* = 0.70, *p* = .330,  *d* = 0.175 |
| 10.94 + cpi | ***t_(15)_* = 3.84, *p* = .003,**  ***d* = 0.960** | ***t_(15)_* = 6.25, *p* = 1.1*10^-5^,**  ***d* = 1.563** | ***t_(15)_* = 3.24, *p* = .003,**  ***d* = 0.811** | ***t_(15)_* = 2.93, *p* = .011,**  *d* = 0.731 |
| Emotion | 0-200 ms | 200-400 ms | 400-600 ms | 600-800 ms |
| 4 cpi | *t_(15)_* = 0.11, *p* = .611,  *d* = 0.027 | *t_(15)_* = 1.31, *p* = .139,  *d* = 0.329 | *t_(15)_* = 0.70, *p* = .493,  *d* = 0.176 | *t_(15)_* = 2.30, *p* = .072,  *d* = 0.575 |
| 4.89 & 5.98 cpi | *t_(15)_* = -1.09, *p* = .853,  *d* = -0.272 | ***t_(15)_* = 2.46, *p* = .027,**  ***d* = 0.614** | *t_(15)_* = 1.61, *p* = .154,  *d* = 0.402 | *t_(15)_* = -0.63, *p* = .732,  *d* = -0.159 |
| 7.31 & 8.94 cpi | *t_(15)_* = 2.15, *p* = .096,  *d* = 0.538 | ***t_(15)_* = 3.41, *p* = .005,**  ***d* = 0.852** | ***t_(15)_* = 1.89, *p* = .047,**  ***d* = 0.472** | *t_(15)_* = 0.98, *p* = .330,  *d* = 0.246 |
| 10.94 + cpi | ***t_(15)_* = 3.45, *p* = .004,**  ***d* = 0.863** | ***t_(15)_* = 6.28, *p* = 1.1*10^-5^,**  *d* = 1.570 | ***t_(15)_* = 3.51, *p* = .002,**  ***d* = 0.877** | ***t_(15)_* = 2.82, *p* = .011,**  ***d* = 0.705** |
| Gender | 0-200 ms | 200-400 ms | 400-600 ms | 600-800 ms |
| 4 cpi | *t_(15)_* = 0.54, *p* = .599,  *d* = 0.134 | *t_(15)_* = 0.78, *p* = .223,  *d* = 0.195 | *t_(15)_* = 1.55, *p* = .283,  *d* = 0.388 | *t_(15)_* = -0.30, *p* = .881,  *d* = -0.074 |
| 4.89 & 5.98 cpi | *t_(15)_* = -0.52, *p* = .853,  *d* = -0.131 | *t_(15)_* = -0.22, *p* = .587,  *d* = -0.056 | *t_(15)_* = 0.07, *p* = .472,  *d* = 0.018 | *t_(15)_* = -0.59, *p* = .732,  *d* = -0.149 |
| 7.31 & 8.94 cpi | *t_(15)_* = 1.21, *p* = .247,  *d* = 0.301 | *t_(15)_* = 1.03, *p* = .159,  *d* = 0.258 | ***t_(15)_* = 2.29, *p* = .037,**  ***d* = 0.572** | *t_(15)_* = 1.21, *p* = .330,  *d* = 0.303 |
| 10.94 + cpi | ***t_(15)_* = 1.76, *p* = .049,**  ***d* = 0.441** | ***t_(15)_* = 2.24, *p* = .020,**  ***d* = 0.561** | ***t_(15)_* = 3.92, *p* = .001,**  ***d* = 0.980** | *t_(15)_* = 0.03, *p* = .489,  *d* = 0.007 |
| Race | 0-200 ms | 200-400 ms | 400-600 ms | 600-800 ms |
| 4 cpi | *t_(15)_* = -1.05, *p* = .846,  *d* = -0.264 | *t_(15)_* = 2.24, *p* = .082,  *d* = 0.559 | *t_(15)_* = -0.25, *p* = .795,  *d* = -0.062 | *t_(15)_* = -0.55, *p* = .881,  *d* = -0.138 |
| 4.89 & 5.98 cpi | *t_(15)_* = 2.19, *p* = .089,  *d* = 0.548 | ***t_(15)_* = 3.66, *p* = .005,**  ***d* = 0.914** | *t_(15)_* = 0.53, *p* = .403,  *d* = 0.132 | *t_(15)_* = 1.09, *p* = .590,  *d* = 0.271 |
| 7.31 & 8.94 cpi | *t_(15)_* = -0.05, *p* = .519,  *d* = -0.012 | ***t_(15)_* = 3.29, *p* = .005,**  ***d* = 0.823** | ***t_(15)_* = 1.79, *p* = .047,**  ***d* = 0.447** | *t_(15)_* = -0.03, *p* = .510,  *d* = -0.007 |
| 10.94 + cpi | ***t_(15)_* = 3.13, *p* = .005,**  ***d* = 0.783** | ***t_(15)_* = 6.24, *p* = 1.1*10^-5^,**  ***d* = 1.560** | ***t_(15)_* = 5.01, *p* = 3.1*10^-4^,**  ***d* = 1.252** | ***t_(15)_* = 2.69, *p* = .011,**  ***d* = 0.672** |

**3. Group-level analysis from Experiment 2-1: temporal sweep SSVEP task**

In this pilot experiment, we investigated face categorization as a function of image presentation duration. With the temporal sweep SSVEP paradigm, images were presented at frequencies of 12, 10, 8, 6, 5, 4, and 3 Hz, following a fast-to-slow presentation mode in each 84-s sequence. We applied identical data processing and analysis methods to our main experiment 2-2.

In the frequency domain analysis, we ran non-parametric bootstrap analysis in the right OT ROI for each face condition, and found significant categorization responses for all four face conditions even when faces were presented at the fastest 12 Hz (i.e., 83 ms/image) (**Figure S2 A**). Scalp topographies (**Figure S2 B**) revealed robust activation patterns in the lateral occipitotemporal (OT) channels, especially in the right hemisphere.

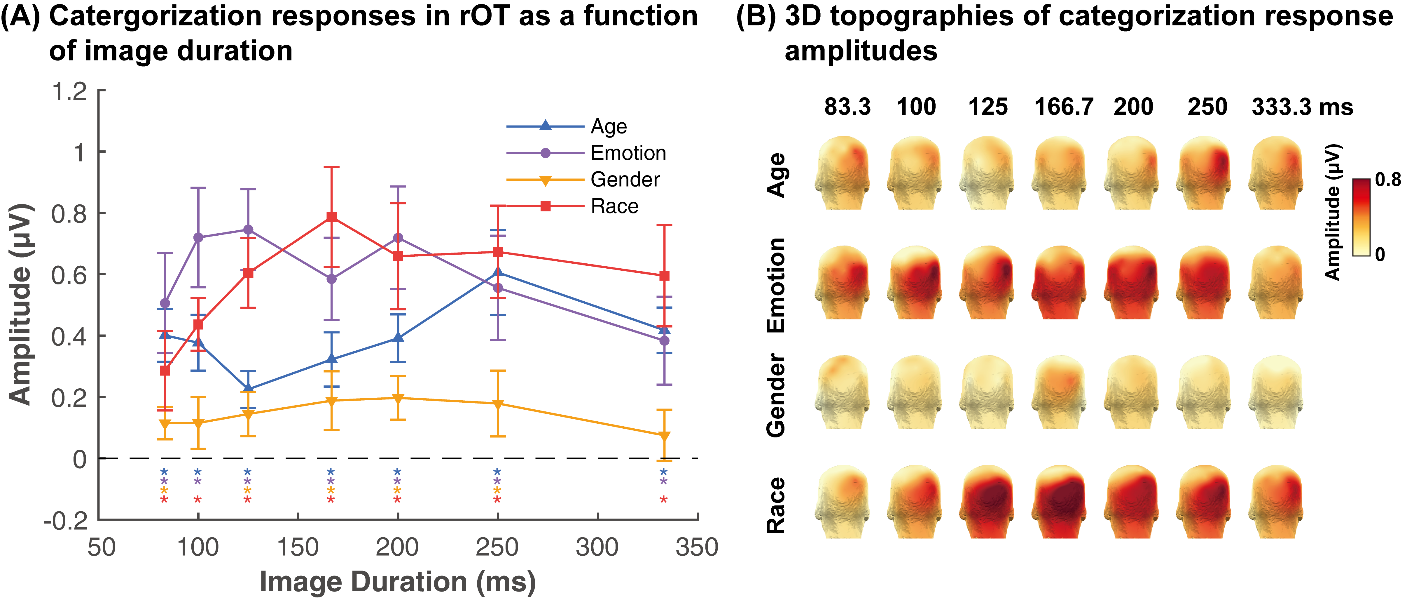

**Figure S2. Face categorization response profiles as a function of image duration for all conditions. A.** Response amplitudes in the rOT ROI. Errors: 95% confidence interval. Asterisks: significant categorization responses from zero. **B.** Three-D topographies.

**Figure S3** presents the time domain results over 17 posterior channels across four face conditions. Results clearly showed consistent spatiotemporal dynamics within conditions but distinct patterns between them. Significant time windows defined with cluster-based permutation tests found significant time windows mainly in 200-600 ms after stimulus onset. Notably, despite using identical target stimuli (young Asian neutral female faces) across all conditions (except emotion, in which sad faces were used), we observed condition-specific spatiotemporal patterns with the SSVEP paradigm, which measured the online discrimination responses between two categories (e.g., young Asian vs. young Caucasian neutral faces). Another interesting finding was that peak response latencies systematically increased with longer image presentation times, particularly for emotion and race processing. This latency shift explains why Experiment 2-2 (brief exposures) showed earlier decoding windows (0–400 ms) compared to Experiment 1 (200 ms exposure; decoding in 0–600 ms), highlighting how temporal constraints dynamically modulate facial feature extraction.

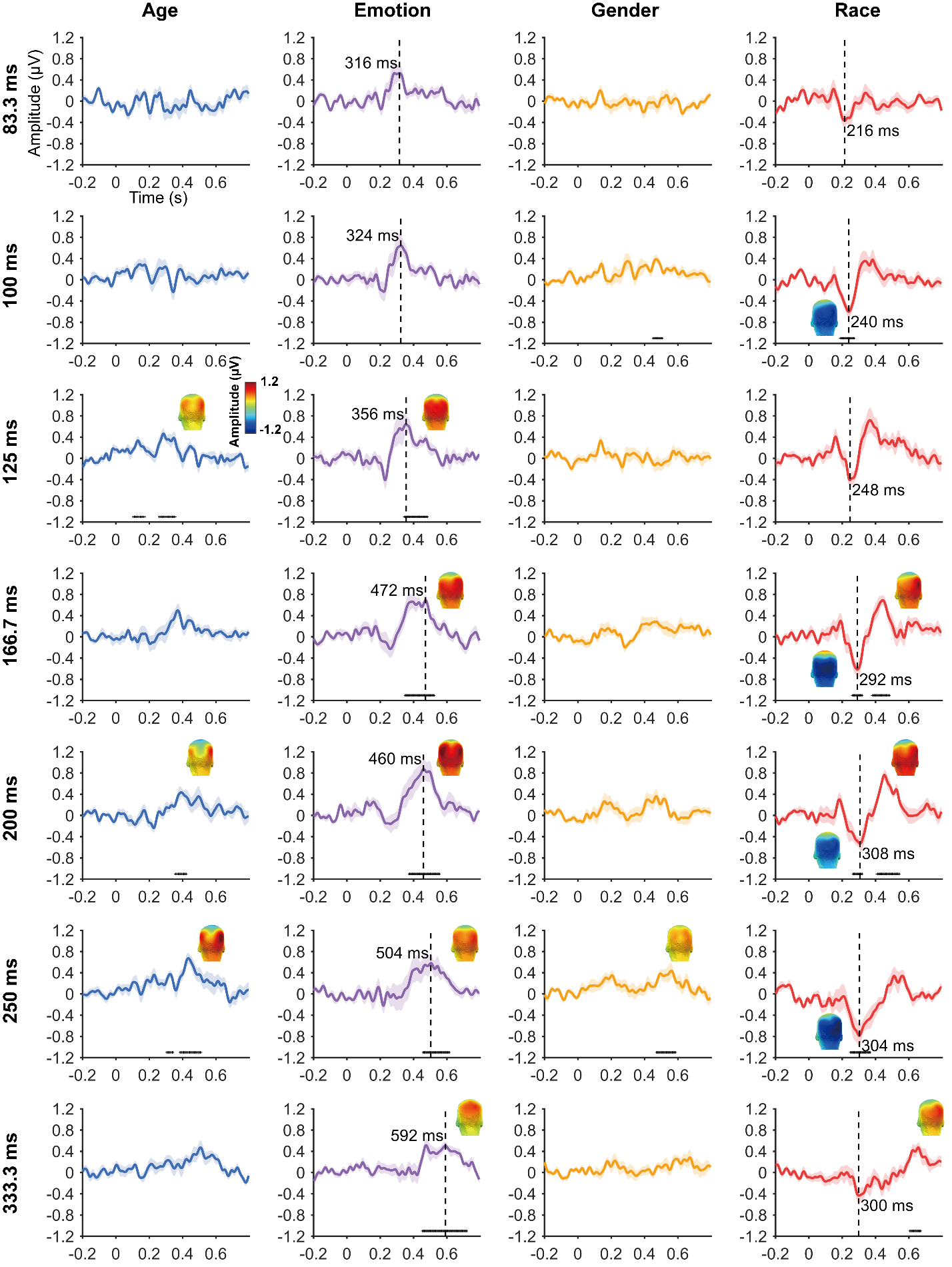

**Figure S3. Temporal dynamics of category-selective responses as a function of image duration.** Data are averaged across channels of a larger posterior ROI and across individuals across different TF steps. Colored lines: mean responses. Shadings: standard errors across participants. Colored horizontal lines at y = -1.1: significant responses relative to zero for the posterior ROI (*ps* < .05). 3D topographies: spatial distribution of the face categorization responses at significant time windows [peak latency $\pm$ 10 ms]. Dashed vertical lines in emotion and race conditions: indicating peak response latencies.

**4. Frequency domain analysis from Experiment 2-2: temporal sweep SSVEP task**

**Figure S4** shows the averaged frequency spectra of face categorization responses at 1 Hz (and its first 8 harmonics) averaged across four face conditions at each TF step over the OT ROI. Significant harmonics (above EEG noise) emerged at step 2, when images were presented at 50 ms. Three-D topographies showed robust activation patterns in the posterior brain regions, especially in the lateral occipitotemporal cortex.

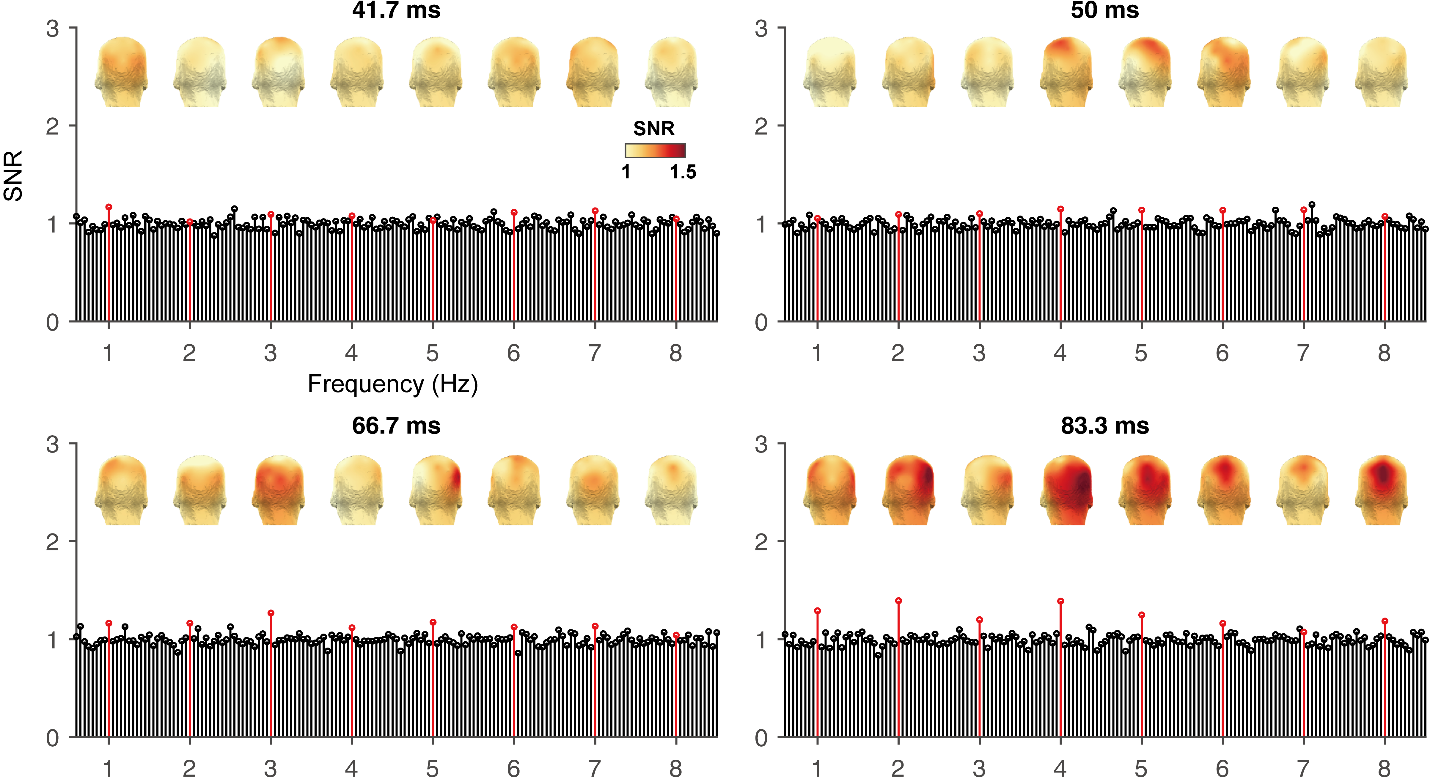

**Figure S4.** **Grand-averaged frequency spectra in signal-to-noise ratio (SNR) across four face conditions as a function of TF steps over the OT ROI.** The frequency bins at every 1 Hz step (up to 8 Hz) are highlighted in red dashed lines. Three-D topographies of the first 8 harmonics of 1 Hz are shown above their corresponding harmonics.

**Figure S5** shows the time domain neural responses across 4 face conditions within an extended posterior ROI. We averaged time courses from the final two temporal frequency (TF) steps, as frequency domain analyses revealed robust categorization responses in all face dimensions at these two steps. Significant time windows for the categorization responses were found in 150-400 ms.

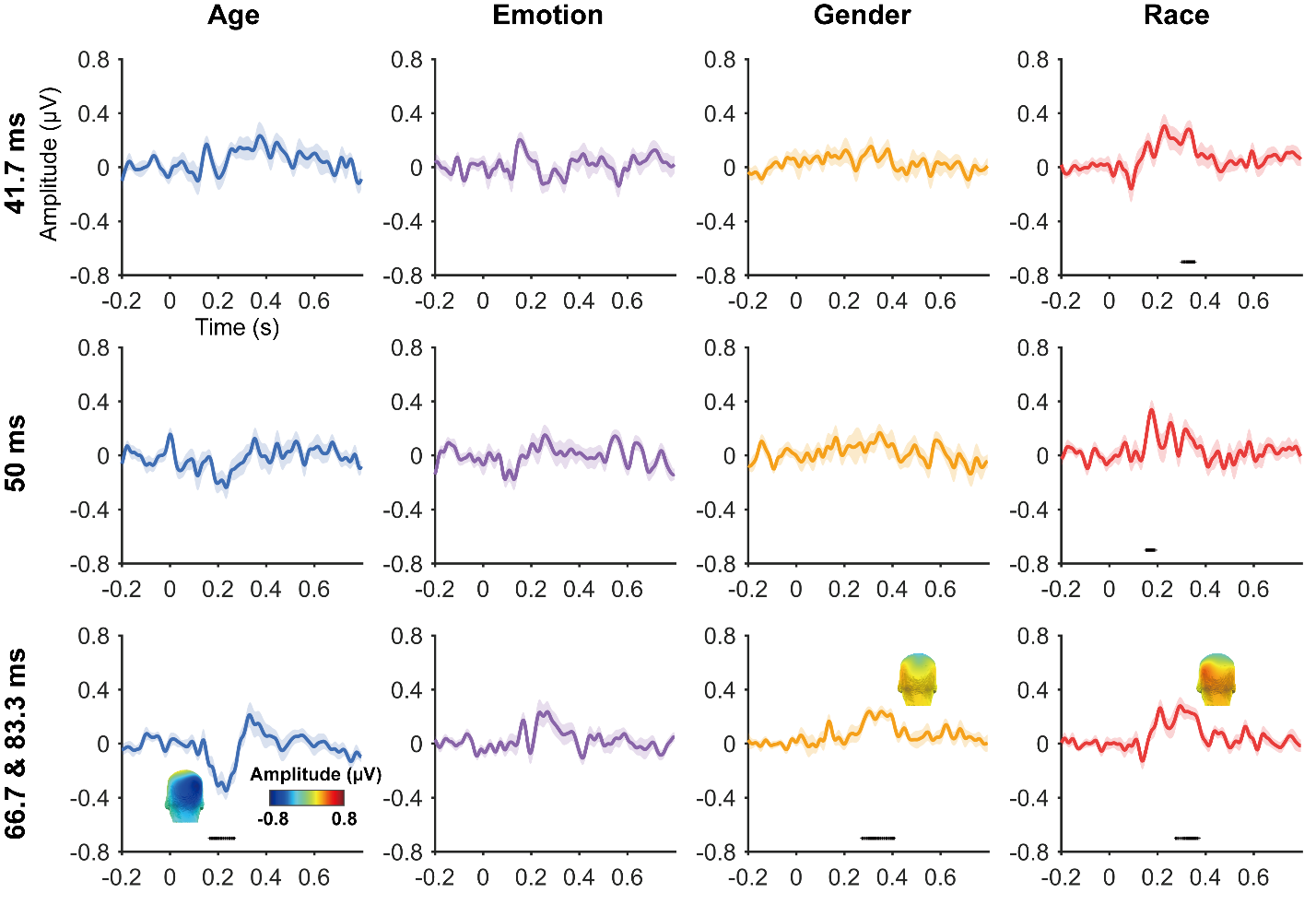

**Figure S5.** **Temporal dynamics of category-selective responses as a function of image duration.** Data are averaged across electrodes of a larger posterior ROI and across individuals. Colored lines: mean responses. Errors: standard errors across participants. Colored horizontal lines at y = -0.7: significant responses relative to zero over the posterior ROI. 3D topographies: spatial distribution of the categorization responses in significant time windows [peak latency $\pm$ 10 ms].

**Table S3. Statistical results for on-diagonal within-condition correlations between odd/even trials in the RSMs across 4 time windows in Experiment 2-2.** Bold fonts: significant above-zero within-condition correlations (FDR-corrected).

| Age | 0-200 ms | 200-400 ms | 400-600 ms | 600-800 ms |
| --- | --- | --- | --- | --- |
| 41.7 ms (24 Hz) | *t_(15)_* = 0.17, *p* = .565,  *d* = -0.042 | *t_(15)_* = 1.15, *p* = .171,  *d* = 0.288 | ***t_(15)_* = 2.60, *p* = .040,**  *d* = 0.651 | *t_(15)_* = 1.41, *p* = .178,  *d* = 0.354 |
| 50 ms (20 Hz) | *t_(15)_* = 1.49, *p* = .104,  *d* = 0.374 | *t_(15)_* = 0.22, *p* = .555,  *d* = 0.054 | *t_(15)_* = -0.97, *p* = .825,  *d* = -0.242 | *t_(15)_* = -2.46, *p* = .987,  *d* = -0.614 |
| 66.7 & 83.3 ms  (15 & 12 Hz) | ***t_(15)_* = 4.89, *p* = 8.8*10^-5^,**  ***d* = 1.245** | ***t_(15)_* = 5.86, *p =* 6.3*10^-5^,**  ***d* = 1.464** | *t_(15)_* = 1.61, *p* = .223,  *d* = 0.402 | *t_(15)_* = -0.34, *p* = .629,  *d* = -0.084 |
| Emotion | 0-200 ms | 200-400 ms | 400-600 ms | 600-800 ms |
| 41.7 ms (24 Hz) | *t_(15)_* = 1.91, *p* = .075,  *d* = 0.478 | *t_(15)_* = 0.98, *p* = .171,  *d* = 0.246 | *t_(15)_* = 0.04, *p* = .647,  *d* = 0.009 | *t_(15)_* = -1.63, *p* = .938,  *d* = -0.408 |
| 50 ms (20 Hz) | *t_(15)_* = 1.89, *p* = .079,  *d* = 0.472 | ***t_(15)_* = 2.71, *p* = .019,**  ***d* = 0.678** | *t_(15)_* = -0.30, *p* = .825,  *d* = -0.076 | *t_(15)_* = 0.82, *p* = .709,  *d* = 0.206 |
| 66.7 & 83.3 ms  (15 & 12 Hz) | ***t_(15)_* = 4.95, *p* =8.8*10^-5^,**  ***d* = 1.237** | ***t_(15)_* = 3.85, *p* = .001,**  ***d* = 0.963** | *t_(15)_* = 1.27, *p* = .223,  *d* = 0.318 | *t_(15)_* = 0.15, *p* = .588,  *d* = 0.038 |
| Gender | 0-200 ms | 200-400 ms | 400-600 ms | 600-800 ms |
| 41.7 ms (24 Hz) | *t_(15)_* = 1.45, *p* = .112,  *d* = 0.362 | *t_(15)_* = 1.80, *p* = .092,  *d* = 0.450 | *t_(15)_* = 1.76, *p* = .098,  *d* = 0.441 | *t_(15)_* = 1.82, *p* = .178,  *d* = 0.454 |
| 50 ms (20 Hz) | *t_(15)_* = 0.89, *p* = .193,  *d* = 0.223 | *t_(15)_* = -1.32, *p* = .897,  *d* = -0.331 | *t_(15)_* = -0.68, *p* = .825,  *d* = -0.169 | *t_(15)_* = 0.22, *p* = .709,  *d* = 0.055 |
| 66.7 & 83.3 ms  (15 & 12 Hz) | ***t_(15)_* = 5.20, *p* = 8.8*10^-5^,**  ***d* = 1.300** | *t_(15)_* = 1.27, *p* = .111,  *d* = 0.318 | *t_(15)_* = 0.85, *p* = .272,  *d* = 0.213 | *t_(15)_* = 0.77, *p* = .455,  *d* = 0.192 |
| Race | 0-200 ms | 200-400 ms | 400-600 ms | 600-800 ms |
| 41.7 ms (24 Hz) | ***t_(15)_* = 3.24, *p* = .011,**  ***d* = 0.810** | *t_(15)_* = 1.89, *p* = .092,  *d* = 0.472 | *t_(15)_* = -0.54, *p* = .703,  *d* = -0.136 | *t_(15)_* = -0.64, *p* = .938,  *d* = -0.161 |
| 50 ms (20 Hz) | ***t_(15)_* = 3.74, *p* = .004,**  ***d* = 0.936** | ***t_(15)_* = 2.63, *p* = .019,**  ***d* = 0.657** | *t_(15)_* = -0.23, *p* = .825,  *d* = -0.058 | *t_(15)_* = -0.08, *p* = .709,  *d* = -0.020 |
| 66.7 & 83.3 ms  (15 & 12 Hz) | ***t_(15)_* = 5.93, *p* = 5.5*10^-5^,**  ***d* = 1.482** | ***t_(15)_* = 5.25, *p* = 9.7*10^-5^,**  ***d* = 1.313** | *t_(15)_* = -0.35, *p* = .633,  *d* = -0.087 | *t_(15)_* = 1.95, *p* = .140,  *d* = 0.488 |

**Table S4. Statistical results for distinctiveness scores for each face condition across four time windows in Experiment 2-2.** Bold texts: significant within-category vs. between-category similarities (FDR-corrected).

| Age | 0-200 ms | 200-400 ms | 400-600 ms | 600-800 ms |
| --- | --- | --- | --- | --- |
| 41.7 ms (24 Hz) | *t_(15)_* = -0.60, *p* = .722,  *d* = -0.151 | *t_(15)_* = 1.67, *p* = .077,  *d* = 0.418 | *t_(15)_* = 2.35, *p* = .066,  *d* = 0.587 | *t_(15)_* = 1.22, *p* = .242,  *d* = 0.305 |
| 50 ms (20 Hz) | *t_(15)_* = 1.87, *p* = .080,  *d* = 0.469 | *t_(15)_* = -0.21, *p* = .777,  *d* = -0.053 | *t_(15)_* = -0.63, *p* = .731,  *d* = -0.157 | *t_(15)_* = -2.78, *p* = .993,  *d* = -0.694 |
| 66.7 & 83.3 ms  (15 & 12 Hz) | ***t_(15)_* = 4.49, *p* = 2.7*10^-4^,**  ***d* = 1.123** | ***t_(15)_* = 5.15, *p* = 2.3*10^-4^,**  ***d* = 1.288** | *t_(15)_* = 1.52, *p* = .297,  *d* = 0.380 | *t_(15)_* = -0.11, *p* = .709,  *d* = -0.028 |
| Emotion | 0-200 ms | 200-400 ms | 400-600 ms | 600-800 ms |
| 41.7 ms (24 Hz) | *t_(15)_* = 1.63, *p* = .124,  *d* = 0.408 | *t_(15)_* = 1.01, *p* = .165,  *d* = 0.252 | *t_(15)_* = -0.09, *p* = .715,  *d* = -0.023 | *t_(15)_* = -1.39, *p* = .907,  *d* = -0.347 |
| 50 ms (20 Hz) | *t_(15)_* = 1.35, *p* = .132,  *d* = 0.337 | ***t_(15)_* = 2.74, *p* = .031,**  ***d* = 0.685** | *t_(15)_* = -0.35, *p* = .731,  *d* = -0.087 | *t_(15)_* = 0.10, *p* = .993,  *d* = 0.026 |
| 66.7 & 83.3 ms  (15 & 12 Hz) | ***t_(15)_* = 4.38, *p =* 2.7*10^-4^,**  ***d* = 1.094** | ***t_(15)_* = 3.40, *p* = .003,**  ***d* = 0.850** | *t_(15)_* = 1.07, *p* = .297,  *d* = 0.269 | *t_(15)_* = -0.56, *p* = .709,  *d* = -0.141 |
| Gender | 0-200 ms | 200-400 ms | 400-600 ms | 600-800 ms |
| 41.7 ms (24 Hz) | *t_(15)_* = 1.19, *p* = .167,  *d* = 0.299 | *t_(15)_* = 1.76, *p* = .077,  *d* = 0.439 | *t_(15)_* = 1.40, *p* = .182,  *d* = 0.350 | *t_(15)_* = 1.72, *p* = .213,  *d* = 0.429 |
| 50 ms (20 Hz) | *t_(15)_* = 0.39, *p* = .352,  *d* = 0.097 | *t_(15)_* = -1.36, *p* = .904,  *d* = -0.341 | *t_(15)_* = 0.06, *p* = .731,  *d* = 0.014 | *t_(15)_* = -0.22, *p* = .993,  *d* = -0.054 |
| 66.7 & 83.3 ms  (15 & 12 Hz) | ***t_(15)_* = 4.61, *p* = 2.7*10^-4^,**  ***d* = 1.152** | *t_(15)_* = 0.74, *p* = .237,  *d* = 0.184 | *t_(15)_* = 0.78, *p* = .297,  *d* = 0.196 | *t_(15)_* = 0.31, *p* = .709,  *d* = 0.078 |
| Race | 0-200 ms | 200-400 ms | 400-600 ms | 600-800 ms |
| 41.7 ms (24 Hz) | ***t_(15)_* = 3.44, *p* = .007,**  ***d* = 0.860** | *t_(15)_* = 2.20, *p* = .077,  *d* = 0.549 | *t_(15)_* = -0.59, *p* = .717,  *d* = -0.147 | *t_(15)_* = -0.84, *p* = .907,  *d* = -0.209 |
| 50 ms (20 Hz) | ***t_(15)_* = 3.66, *p* = .005,**  ***d* = 0.916** | *t_(15)_* = 2.11, *p* = .053,  *d* = 0.526 | *t_(15)_* = -0.07, *p* = .731,  *d* = -0.019 | *t_(15)_* = -0.67, *p* = .993,  *d* = -0.169 |
| 66.7 & 83.3 ms  (15 & 12 Hz) | ***t_(15)_* = 4.92, *p* = 2.7*10^-4^,**  ***d* = 1.231** | ***t_(15)_* = 4.34, *p* = 5.8*10^-4^,**  ***d* = 1.085** | *t_(15)_* = -0.64, *p* = .733,  *d* = -0.159 | *t_(15)_* = 1.65, *p* = .241,  *d* = 0.412 |
